# TNF Blockade Reduces Prostatic Hyperplasia and Inflammation while Limiting BPH Diagnosis in Patients with Autoimmune Disease

**DOI:** 10.1101/2021.03.11.434972

**Authors:** Renee E. Vickman, LaTayia Aaron-Brooks, Renyuan Zhang, Nadia A. Lanman, Brittany Lapin, Victoria Gil, Max Greenberg, Takeshi Sasaki, Gregory M. Cresswell, Meaghan M. Broman, Jacqueline Petkewicz, Pooja Talaty, Brian T. Helfand, Alexander P. Glaser, Chi-Hsiung Wang, Omar E. Franco, Timothy L. Ratliff, Kent L. Nastiuk, Susan E. Crawford, Simon W. Hayward

## Abstract

Benign prostatic hyperplasia (BPH) is ostensibly linked to autoimmune (AI) diseases, but whether the prostate is a target of systemic inflammation associated with AI conditions is unknown. Prostatic inflammation is linked to fibrosis, hyperplasia, and reduced responses to BPH-related medical therapies. This study was conducted to determine if AI disease correlates with BPH diagnosis and whether systemic targeting of an inflammatory mediator limits prostatic inflammation and hyperplasia. Patient medical records (n=112,152) were evaluated to determine BPH prevalence among different AI diseases. Inflammatory cells from human BPH tissues were analyzed by single-cell (sc)RNA-seq and the tumor necrosis factor (TNF)α-antagonist etanercept was tested in two murine models of prostatic enlargement. BPH prevalence was significantly higher among patients with AI disease compared to unaffected individuals. However, AI patients treated with TNFα-antagonists had a significantly reduced incidence of BPH. Data from scRNA-seq identified macrophages as a dominant source of TNFα and *in vitro* assays confirmed that TNFα stimulates BPH-derived fibroblast proliferation. In the AI patient cohort and murine models, systemic treatment with TNFα-antagonists decreased prostatic epithelial proliferation, macrophage infiltration, and epithelial NFκB activation compared to control tissues. These studies are the first to show that patients with AI diseases have a heightened susceptibility to BPH and that the TNFα-signaling axis is important for BPH pathogenesis. Macrophage-secreted TNFα may mechanistically drive BPH via chronic activation of the signaling axis and NFκB. TNFα blockade appears to be a promising new pharmacological approach to target inflammation and suppress BPH.

**One sentence summary:** Patient data and mouse models suggest that repurposing tumor necrosis factor alpha blockade reduces inflammation-mediated prostatic hyperplasia.

## Introduction

The vast majority of men develop histological benign prostatic hyperplasia (BPH) and nearly half exhibit moderate to severe clinical symptoms before age 80 (*1, 2*). The precise mechanisms underlying the pathogenesis of BPH and how it contributes to lower urinary tract symptoms (LUTS) are not well understood. The only definitive risk factors for developing BPH are male sex and increasing age, but BPH has been linked to decreased systemic androgen:estrogen ratios, obesity, type 2 diabetes, metabolic syndrome, and inflammation (*3–9*). There is a paucity of studies investigating possible genetic determinants of BPH, although, more recently, there has been greater interest in this area (*10–12*). The medical therapeutic options used for men with LUTS related to BPH are limited (e.g. alpha-adrenergic antagonists and 5α-reductase inhibitors [5ARIs] (*13*)) and have not changed significantly for two decades. Due to therapeutic resistance or disease progression, over 100,000 men undergo surgical procedures for BPH each year in the United States (*14*). A nonsurgical, targeted approach for medical treatment of these BPH cases is needed.

Chronic inflammation in the prostate can contribute to prostatic hyperplasia, fibrosis, and failure to respond to therapy in BPH (*13, 15–17*). CD45+ leukocytes are known to comprise a significant percentage of cells in BPH tissues, with macrophages and T cells as major populations (*18*). CD68+ macrophages accumulate in BPH tissues and aid in stromal cell proliferation (*19*). Whether BPH is a component of a systemic inflammatory process is not known. BPH has been suggested to have characteristics of autoimmune (AI) inflammatory conditions (*20, 21*), consistent with our observations of an inflammatory gene expression signature including activation of AP-1 stress factors associated with severely symptomatic BPH that was refractory to medical therapy and required surgical intervention (*22*). BPH is associated with a number of common pro-inflammatory comorbidities (*4–8*), suggesting that a systemic pro-inflammatory environment may promote hyperplasia and/or exacerbate symptoms. Although there is no perfect animal model to study human BPH, some, such as non-obese diabetic (NOD) mice, recapitulate the association of chronic prostatic inflammation with prostatic hyperplasia, similar to the observations in human disease (*23*).

AI conditions such as rheumatoid arthritis (RA) and systemic lupus erythematosus are associated with systemic inflammation and can involve multiple organ sites; however, the prostate has not been recognized as a target organ of these inflammatory processes. Interestingly, AI diseases have similar co-morbidities to BPH, including obesity, type 2 diabetes, and metabolic syndrome (*24–27*), and some AI conditions are also comorbidities of other AI diseases, such as psoriasis and inflammatory bowel disease (*28*). The earliest and most widely used therapeutics for targeting a specific biological pathway in AI conditions are tumor necrosis factor (TNF)α- antagonists, which limit the inflammatory properties of this cytokine (*29*). AI diseases are significantly more prevalent in women compared to men, potentially highlighting an immunosuppressive role of androgens in protecting against AI diseases (*30–32*). Recent single-cell mRNA-sequencing (scRNA-seq) studies in AI diseases have identified a variety of immune cell populations including monocytes/macrophages, T cells, and B cells in diseased tissues (*33–35*). Notably, ligand-receptor pair interaction analyses highlight the impact of macrophage-secreted TNFα on stromal cells in Crohn’s disease (*35*). It is clear that TNFα ligand-receptor signaling is important within AI diseases, but the contribution of this cytokine to the development or progression of BPH in this patient population has not been studied.

To understand the underlying processes which drive BPH pathogenesis, we investigated whether BPH prevalence is increased among men with AI conditions and tested whether TNFα- antagonists suppress prostatic inflammation in BPH. Patient medical records and human prostate tissues were used to support repurposing of approved therapeutics. The results suggest that TNFα- antagonists are a valid therapeutic for reducing BPH incidence and decreasing localized inflammation within the prostate.

## Results

### BPH prevalence is elevated in specific autoimmune inflammatory conditions

This study received institutional review board (IRB) approval. A retrospective evaluation of the NorthShore University HealthSystem Enterprise Data Warehouse was conducted to determine whether patients with autoimmune (AI) disease had an elevated risk of BPH diagnosis. Male patients over the age of 40 who had a NorthShore office visit between 01/01/2010- 12/31/2012 were included (n=112,152). Patients with a diagnosis of prostate cancer were excluded from evaluation. Records were searched for diagnoses of BPH and a range of AI conditions, most commonly psoriasis, RA, and ulcerative colitis. The cohort included 101,383 (90.4%) men with no history of AI and 10,769 (9.6%) men with a diagnosis of one or more AI conditions (Figure 1A). To determine whether treatment for AI diseases affected BPH diagnosis in this population, patients with an AI disease diagnosis were further divided into those who had a diagnosis of AI disease *prior to* a BPH diagnosis or those who had a diagnosis of AI disease *after* a BPH diagnosis (Figure 1A). Results from a chi-square test indicate that the baseline prevalence of BPH was 20.3% in patients with no history of AI disease, but patients with diagnosed AI conditions had a significantly increased BPH prevalence of 30.6% (p<0.001; Figure 1B-C; Table S1). The most marked increases in BPH prevalence were associated with RA (38.0%), type 1 diabetes (32.0%), lupus (30.7%), ulcerative colitis (30.2%), and Crohn’s disease (27.4%), while other diseases, such as multiple sclerosis (21.6%), showed little change from baseline (Figure 1D; Table S1).

**Figure 1.**
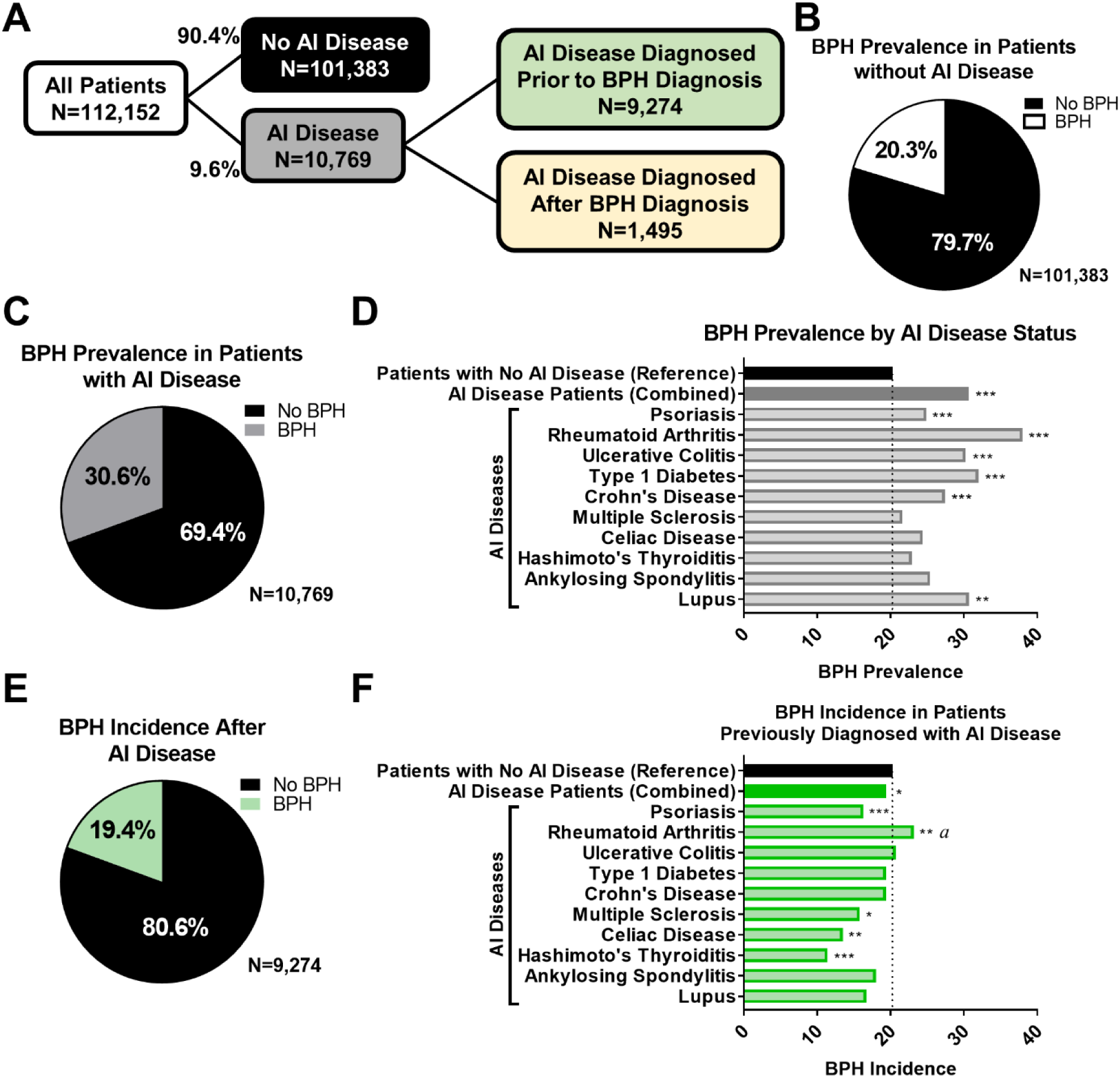
Men with autoimmune disease have increased BPH prevalence. Chi-square tests were utilized to compare the proportion of BPH diagnoses in men with an AI condition *versus* men with no AI condition. **A)** Flow chart indicating the breakdown of patients into groups based on the presence of AI disease diagnosis (9.6% with and 90.4% without AI disease diagnosis). Patients with AI disease diagnosis were further separated into groups based on whether AI disease diagnosis occurred prior to or after BPH diagnosis. **B)** BPH prevalence in patients without AI disease is 20.3%. **C)** BPH prevalence in patients with AI disease is 30.6%. **D)** Graph indicates the significant increase in BPH prevalence in patients with different AI diseases compared to patients without AI disease. **E)** The BPH incidence in patients previously diagnosed with AI conditions, when patients may have been treated for these conditions, is 19.4%. **F)** Graph indicates the significant changes in BPH incidence in patients diagnosed with different AI diseases *prior to* BPH diagnosis compared to the baseline BPH prevalence of 20.3%. *^a^* indicates a significantly higher BPH incidence than the 20.3% reference, although this is decreased from 38.0% prevalence in all RA patients (D).

**Table 1.**
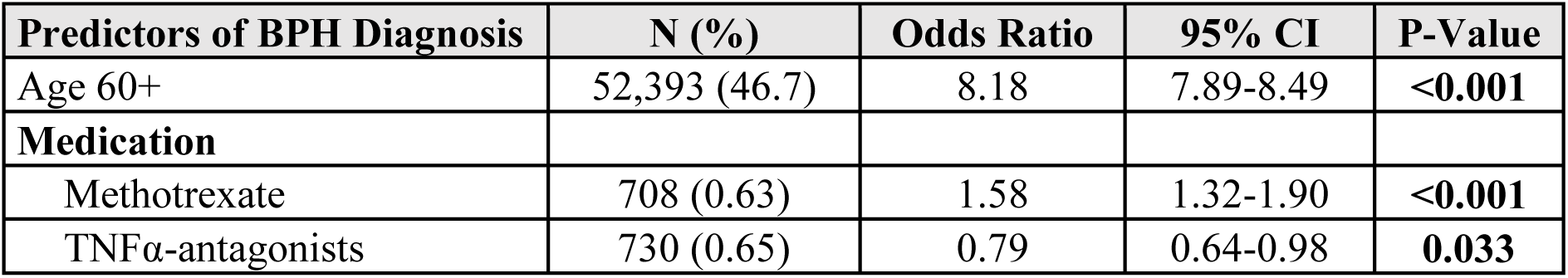
Modeling predictors of BPH diagnosis indicate that treatment of patients with TNF- antagonists is protective for BPH. Multivariable logistic regression models were constructed to identify predictors of BPH diagnosis. Predictors were determined *a priori*: age over 60 years, methotrexate, and TNFα-antagonists. These studies included all 112,152 patients in the cohort.

| Predictors of BPH Diagnosis | N (%) | Odds Ratio | 95% CI | P-Value |
| --- | --- | --- | --- | --- |
| Age 60+ | 52,393 (46.7) | 8.18 | 7.89-8.49 | <0.001 |
| **Medication** |  |  |  |  |
| Methotrexate | 708 (0.63) | 1.58 | 1.32-1.90 | <0.001 |
| TNFα-antagonists | 730 (0.65) | 0.79 | 0.64-0.98 | **0.033** |

In the subset of men who were diagnosed with their AI condition *prior to* BPH diagnosis, the BPH incidence was 19.4%, similar to the baseline BPH prevalence (p=0.037; Figure 1E; Table S1). This suggests that treatment of AI disorders significantly diminishes subsequent BPH diagnoses, although there were some disease-specific variations in BPH incidence with treatment (Figure 1F; Table S1). Nearly all AI conditions in this group had a significantly lower incidence of BPH compared to the baseline in patients without AI disease, with the exception of RA patients who remained at a significantly elevated rate of BPH diagnosis compared to control patients (23.2%; p=0.007). However, this BPH incidence was reduced from the BPH prevalence in all RA patients (38.0%; Figure 1D, F).

### Patients treated with TNFα-antagonists for autoimmune conditions have decreased BPH diagnoses

Using the complete patient population from Figure 1, patient age and the use of common therapeutics, specifically methotrexate and TNFα-antagonists, were modeled as predictors of BPH diagnosis using multivariable logistic regression. Subjects over the age of 60 were significantly more likely (odds ratio=8.18, p<0.001) to have BPH than those under age 60 (Table 1). This analysis indicated that treatment with specific therapeutics, namely TNFα-antagonists but not methotrexate, significantly decreased (odds ratio=0.79, p=0.033) the likelihood of a BPH diagnosis compared to patients not taking these drugs (Table 1). This effect was independent of age. Further, the median time for any BPH patient that progressed to surgery from the time of diagnosis was 616 days (n=1,685). The median time for BPH patients to progress from diagnosis to BPH-specific surgeries was 728 days in individuals receiving methotrexate (n=15, p=0.75) *versus* no methotrexate or 1,115 days in patients taking TNFα-antagonist therapy (n=7, p=0.15) *versus* no TNFα-antagonists (Table S2). The increase in the time of progression to BPH-related surgeries for patients taking TNFα-antagonists did not reach significance due to the smaller number of treated patients who progressed to surgery, although this is suggestive that TNFα- antagonist treatment may prolong the time course of BPH patients toward a surgical endpoint.

### scRNA-seq of CD45+ cells from human BPH tissues implicate T cells and macrophages as key sources of TNF

The decrease in BPH incidence after treatment with systemic anti-inflammatory agents that target TNFα suggests a role for inflammation in BPH pathogenesis. Therefore, scRNA-seq studies were pursued to characterize inflammatory cells within prostate tissues from patients with informed consent, per the IRB-approved NorthShore Tissue Biorepository. As prostate size increases with BPH, immune cell density also increases (p=0.0021; Figure 2A-B). To evaluate the TNFα-producing and TNFα-responding inflammatory cell types within human BPH tissues, we conducted scRNA-seq analysis on CD45+ cells isolated from two groups. The first group represented limited prostatic growth and included cells from the transition zone of smaller (<40 grams, n=10) prostates, whereas the second group represented prostatic enlargement using large (>90 grams, n=4) prostates (Figures 2C; S1). Prostate tissues were isolated from age-matched (p=0.32) and body mass index (BMI)-matched (p=0.13) patients (Figure S2A-B). Patients with larger prostates exhibited significantly higher International Prostate Symptom Scores (IPSS) compared to patients with smaller prostates (p=0.0007; Figure S2C-D).

**Figure 2.**
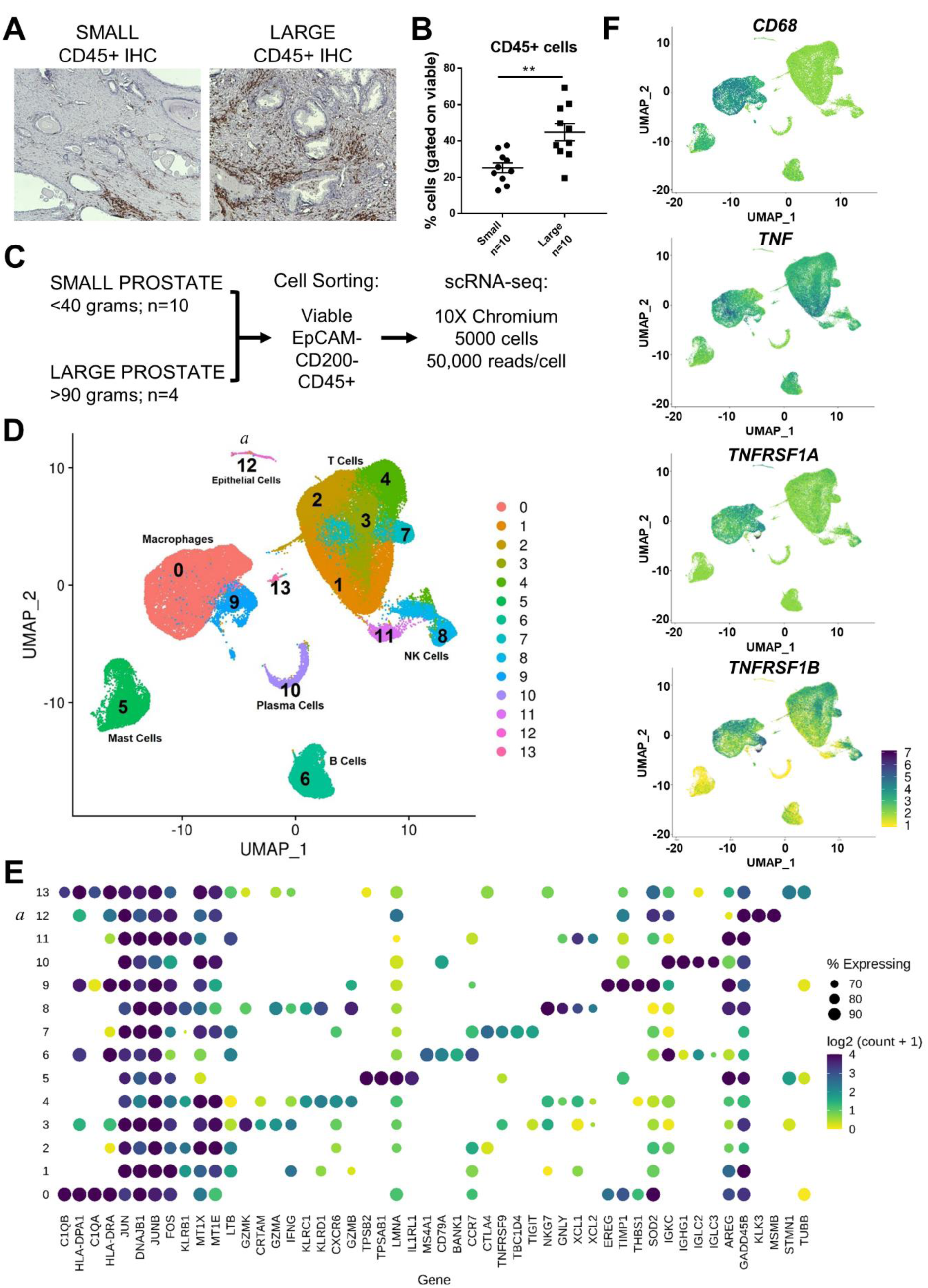
Analysis of CD45+ cells from human BPH tissues indicate macrophages express high levels of TNF and TNF receptors. **A)** CD45 IHC in human prostate transition zone from small or large prostates. **B)** Human prostate transition zone from small or large prostates were digested, stained, and analyzed by flow cytometry. Graph indicates % CD45+ cells, gated on viable cells. **C)** Schematic representing the setup for scRNA-seq studies of human BPH leukocytes (CD45+ cells). A total of 10 small and 4 large prostate transition zone tissues were digested and CD45+EpCAM-CD200- cells sorted by FACS (*72*). scRNA-seq was conducted using the 10X Chromium system, aiming for 5000 cells/sample at a depth of 50,000 reads/cell. **D-F)** scRNA-seq of BPH associated leukocytes. **D)** Uniform manifold approximation and projection (UMAP) plot of 69,850 individual cells, demonstrating dominant T cell and macrophage populations. **E)** Dot plot of the top 4 marker genes from each cluster ranked by fold-change. Gene names are shown on the x-axis and clusters on the y-axis. The size of the dots corresponds to the percentage of cells in a given cluster that express the marker gene. The color of the dots represents the mean log2(counts+1) of each gene in the corresponding cluster. **F)** Feature plots highlighting gene expression of *CD68*, *TNF*, *TNFRSF1A* (TNFR1), and *TNFRSF1B* (TNFR2). *^a^* indicates a small contaminating epithelial population as cluster 12.

Unsupervised clustering analysis of scRNA-seq data allowed for the identification of various cell populations using differential gene expression and cell surface protein expression using CITE-seq (*36*). T cells (CD3+ and CD4+ or CD8+) and macrophages (CD11b+) comprised the dominant immune cell populations but B cells (CD19+), mast cells, NK cells, and plasma cells were also identified (Figures 2D-E; S2E). Although no significant differences in overall immune subpopulations were noted when comparing large *versus* small tissues (Figure S2F), several of the most significantly altered pathways between large and small samples included pathways related to AI conditions (Table S3). The percentage of CD11b+ myeloid cells, CD19+ B cells, and CD4+ or CD8+ T cells from the original digested samples by flow cytometry analysis were correlated with the percentage of these cell types based on cluster identification in the scRNA-seq analysis. Linear regression identified significant correlations for myeloid cells, B cells, and CD8+ T cells, indicating that the cell types identified by scRNA-seq were representative of the CD45+ population in the digested human tissues (Figure S3). Cells with the highest expression of *TNF* were within the T cell and macrophage compartments, and macrophages (highlighted as CD68+) also expressed high levels of TNFα receptors 1 and 2, *TNFRSF1A* and *TNFRSF1B*, respectively (Figures 2F, S4). Macrophage cluster 0 had significantly higher expression of *TNF* (fold-change=1.364, p<0.0001) and *TNFRSF1A* (fold-change=1.325, p<0.0001), while cluster 9 had significantly higher expression of *TNFRSF1A* (fold-change=1.706, p<0.0001), compared to all other clusters (Figure S4). Since BPH macrophages had elevated expression of genes for both expression and response to TNFα, we focused on these cells.

### TNFα-antagonist treatment causes prostatic regression and reduces prostatic epithelial NFκB in Pb-PRL mice

To assess the impact of TNFα-antagonists on BPH and prostatic inflammation, two mouse models of prostatic enlargement were utilized: a transgenic mouse model with prostate-specific expression of the hormone prolactin (probasin-prolactin [Pb-PRL]) that exhibits prostatic enlargement associated with extensive interstitial inflammation (*37, 38*) as well as a spontaneous model of autoimmune inflammation-associated prostatic hyperplastic growth (the NOD mouse) (*23*).

Pb-PRL mice have been shown to develop prostatic hyperplasia with associated interstitial inflammation (*37, 38*). Aged Pb-PRL mice (20-22 months) were treated with 4 mg/kg TNFα- antagonist etanercept or PBS control twice weekly for 12 weeks (n=6 in control group, n=5 in treated group). The dosing regimen of etanercept was sufficient to block intraprostatic TNFα signaling (*39*). Ventral prostate volume was monitored by ultrasound every four weeks using high-resolution, high-frequency ultrasound (*40*). Ventral prostate volume trended downward after eight weeks and was reduced over 30% after 12 weeks of treatment (p=0.0365; Figure 3A). Histological evaluation of prostate tissues following etanercept treatment indicated significantly diminished epithelial cell proliferation via Ki67 immunohistochemistry (IHC) staining compared to control tissues (p=0.032; Figure 3B-C). Evaluation of Pb-PRL prostate tissues of etanercept-treated *versus* control mice did not yield significant differences for either macrophage infiltration as a percentage of immune cells or epithelial NFκB activation via phospho-p65 staining (p=0.782 and p=0.547, respectively; Figure S5).

**Figure 3.**
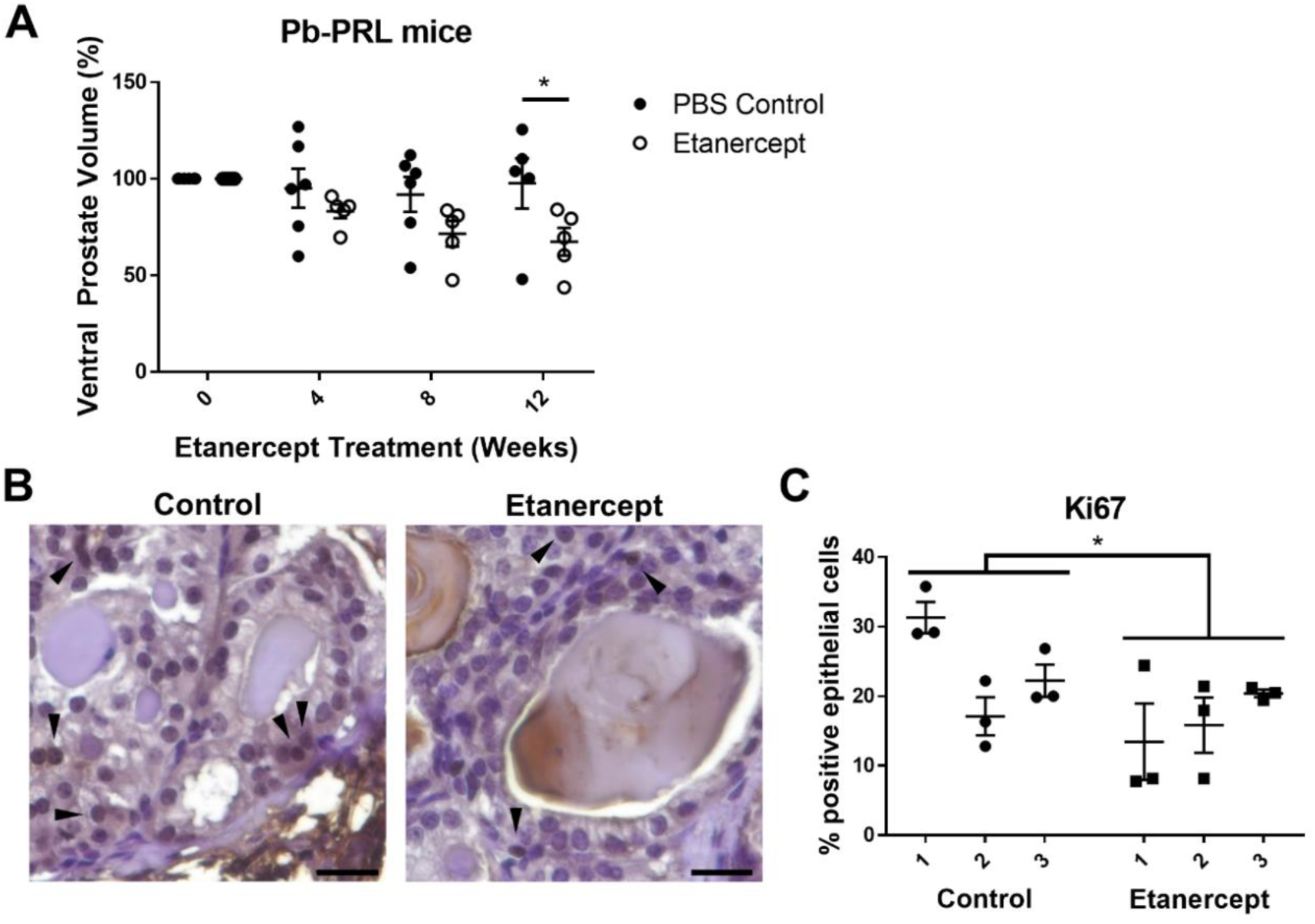
TNFα-antagonist treatment reduces prostate size and epithelial proliferation in Pb- PRL mice. Pb-PRL mice (20-22 months) were treated with 4 mg/kg etanercept or PBS vehicle for 12 weeks. **A)** Volume of ventral prostate by ultrasound every four weeks during the 12-week treatment with etanercept or PBS vehicle control. Measurements are normalized to pre-treatment volume (151 +/- 10 mm^3^), and the plot indicates the mean +/- SEM. One control mouse was removed from the 12-week evaluation. Linear model analysis revealed a significant difference based on treatment (p<0.0001) and Bonferroni correction determined a reduction in ventral prostate volume compared to PBS-treated mice at 12 weeks (p=0.0365; n=5 per group). **B)** Representative images of Ki67 staining in control or etanercept treated mice. **C)** Quantitation of IHC staining for epithelial Ki67 in control or etanercept treated mice, indicated as the percent Ki67 positive epithelial cells per field of view. Data indicate the mean +/- SEM of the percent positive cells in at least three prostate tissue fields for each animal. Comparison of control and etanercept- treated groups for statistical purposes were conducted using a mixed effects model. Scale bars = 20µm.

### TNFα-antagonist treatment reduces prostate hyperplasia, inflammation, and epithelial NFκB in NOD mice

We recently reported that NOD mice have inflammation-associated prostatic hyperplasia (*23*). Therefore, these mice were used as a model to understand the consequence of TNFα blockade on hyperplastic expansion. Animals were treated with 4 mg/kg etanercept or vehicle control twice weekly for five weeks, starting at 25 weeks, as indicated in Figure 4A. Diabetic status was evaluated at the time of first and last treatment, although our previously published work indicated that regions of inflammation, rather than diabetic status, primarily impacts prostatic hyperplasia in NOD mice (*23*). Therefore, diabetic and non-diabetic animals were pooled for analysis of treatment groups. Quantitation of Ki67 IHC staining indicated a reduction in epithelial proliferation in etanercept-treated *versus* control mice (p=0.005; Figure 4B-C), consistent with the Pb-PRL model. Evaluation of prostatic macrophages via F4/80 staining demonstrated a significant decrease in the percentage of macrophages among total immune cells in etanercept-treated *versus* control mice (p=0.002; Figure 4D-E). Furthermore, etanercept treatment caused a reduction in epithelial NFκB activity, as demonstrated by a significant reduction in epithelial phospho-p65 staining (p<0.0001; Figure 4F-G). These data provide an association between the inflammatory cytokine TNFα with epithelial proliferation and NFκB signaling in the prostate. The differences between the Pb-PRL and NOD models could reflect differences in age at treatment or the nature of the model, since NOD is a spontaneous inflammatory model and Pb-PRL is an androgen-driven transgenic with unknown contributions of inflammatory status on androgenic stimulation.

**Figure 4.**
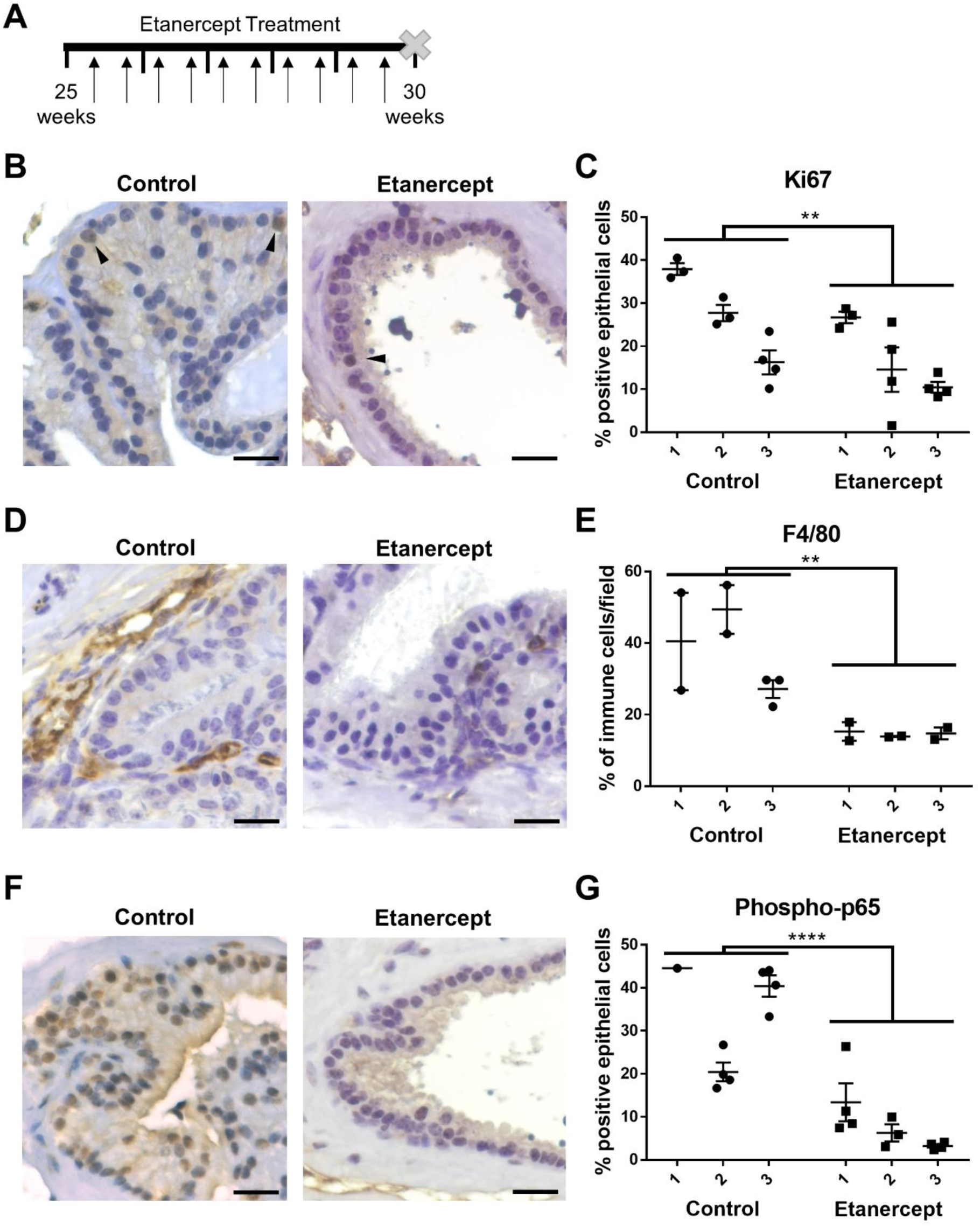
TNFα-antagonist treatment reduces prostatic epithelial proliferation, macrophage infiltration, and NFκB activity in NOD mice. **A)** Diagram representing the timeline of treatment in NOD mice. Mice were treated twice per week for 5 weeks with 4 mg/kg etanercept or PBS vehicle, indicated with black arrows. At the end of the 5-week treatment period, tissues were harvested for analysis. **B)** Representative images of Ki67 staining in control or etanercept-treated mice. **C)** Graph indicating the quantitative summary of Ki76 IHC as the percent of positive epithelial cells per field. **D)** Representative images of F4/80+ staining in control or etanercept-treated mice. **E)** Quantitative summary of IHC staining for F4/80+ cells, represented as the portion of all immune cells in each field. **F)** Representative images of phospho-p65 staining in control or etanercept-treated mice. **G)** Data presented indicate the percentage of phospho-p65 positive epithelial cells counted per field of view. **C, E, G)** Data indicate the mean +/- SEM of the percent positive cells in the indicated number of prostate tissue fields for each animal (n=3 per group). Comparison of control and etanercept-treated groups for statistical purposes were conducted using a mixed effects model and asterisks indicate the significance of the treatment. Scale bars = 20µm.

### TNFα stimulates fibroblast proliferation but not epithelial cell proliferation in vitro

Fibroblasts are an essential component of the prostate microenvironment during organogenesis and in BPH pathogenesis (*41*). Since TNFα-antagonist treatment significantly reduces epithelial proliferation in two independent mouse models, we tested whether TNFα directly influences epithelial and fibroblast cell growth *in vitro*. Importantly, treatment of two human benign prostatic epithelial cell lines elicited no changes in cell growth response to 1 or 10ng/mL TNFα (Figure S6A-B). In contrast, similar treatment with TNFα in a human benign prostatic stromal cell line enhanced cell proliferation (p<0.0001; Figure S6C). TNFα also directly stimulated the proliferation of primary human prostatic fibroblast cultures from two different patients (p<0.0001; Figure 5A). Furthermore, treatment of primary BPH fibroblasts with conditioned medium from M1- or M2-polarized THP-1 cells demonstrated that macrophage-secreted factors stimulate fibroblast growth (Figure 5B). The addition of a TNFα neutralizing antibody indicated that macrophage-stimulated growth can be TNFα-dependent (p=0.0011 and p=0.0148 for M1 and M2, respectively, in patient 376), although this may be patient-specific (Figure 5B). Since epithelial cell proliferation was not directly affected by TNFα treatment *in vitro*, these data are consistent with the long-standing idea that prostate expansion in BPH has a significant stromal input *in vivo*.

**Figure 5.**
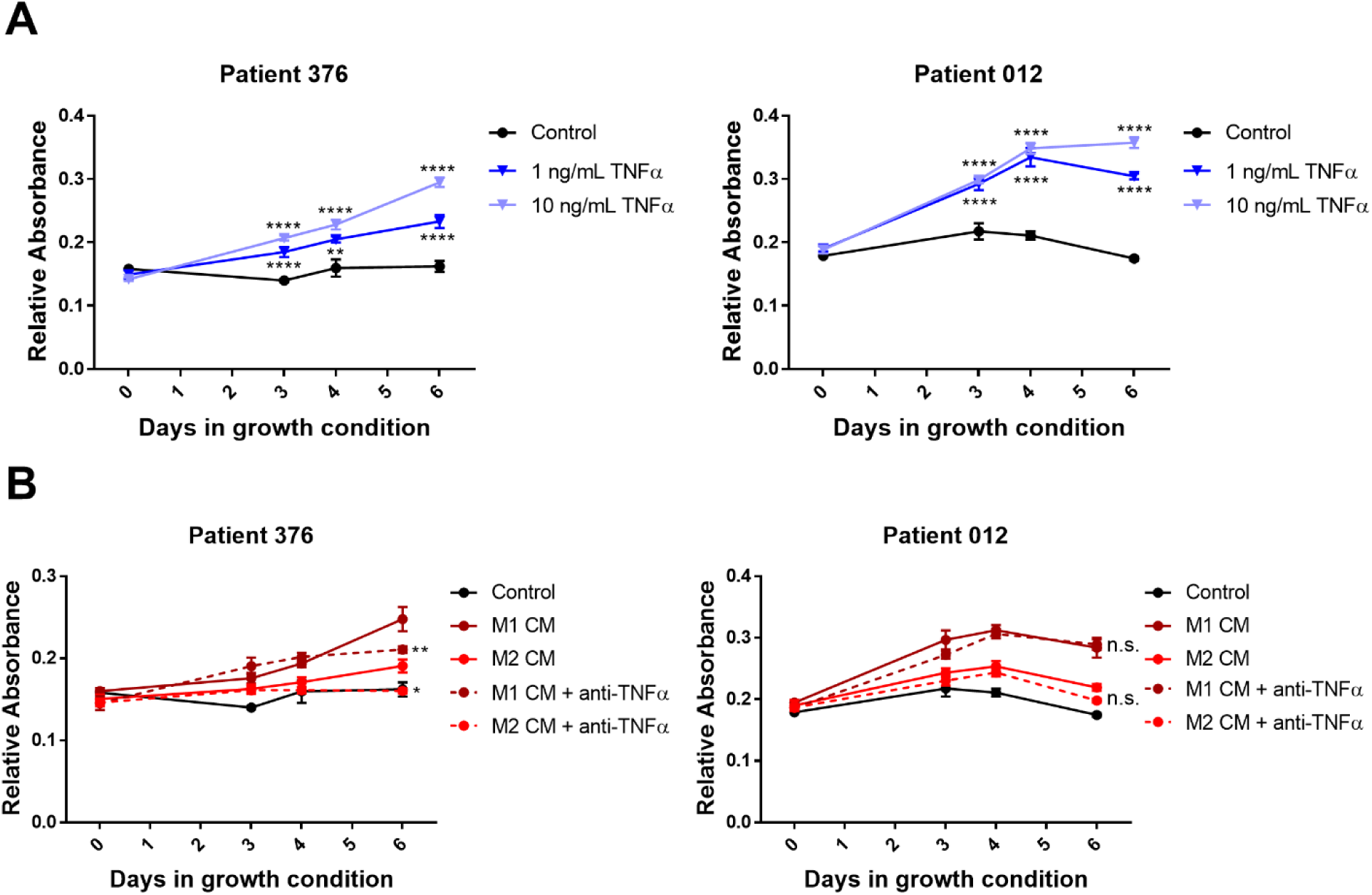
Macrophage-derived TNFα promotes prostate fibroblast cell growth. Primary prostate fibroblasts from two BPH tissues (patients 376 and 012) were subjected to indicated treatments. Crystal violet growth assays were performed in low serum conditions (0.5%) over six days. **A)** Fibroblast cultures were grown in the presence or absence of 1 or 10 ng/mL recombinant TNFα. Asterisks indicate significant differences compared to control samples, determined by two- way ANOVA with multiple comparisons test. **B)** Fibroblast cultures were grown in the presence of 50% M1 or M2 macrophage conditioned medium (generated from THP-1 cells) +/- 40 µg/mL TNFα neutralizing antibody. Points indicate the mean +/- SEM of at least five technical replicates and graphs are representative of three independent experiments. Asterisks indicate significant differences compared to the paired conditioned medium condition without anti-TNFα neutralization, using a two-way ANOVA with post-hoc multiple comparisons test. N.s.=not significant

### TNFα-antagonist treatment in human patients decreases inflammation and epithelial proliferation in the prostate

To better understand the impact of TNFα-antagonist treatment on human prostate tissues, we performed a retrospective study using de-identified human prostate tissues collected through the NorthShore Urologic Disease Biorepository. The biorepository provided transition zone tissues from patients taking TNFα-antagonists who also underwent surgery for prostatic diseases (n=5). Age- and BMI-matched patient samples were used as controls (n=4; Figure S7A-B). All of the patients in both the treated and control groups had a RALP due to cancer, but also had a clinical diagnosis of BPH. No significant changes in prostate volumes or IPSS were observed in treated patients compared to controls (Figure S7C-D). Benign tissue regions from patients treated with TNFα-antagonists had significantly less epithelial Ki67 staining compared to tissues from control patients (p=0.007; Figure 6A-B). In general, the inflammatory process appeared to be lower in tissues from treated patients, and quantitative evaluation of CD68+ cells via IHC indicated that TNFα-antagonist treatment significantly reduced the percentage of CD68+ macrophages out of total immune cells compared to controls (p<0.0001; Figure 6C-D). Tissues from TNFα-antagonist treated patients also had significantly diminished epithelial NFκB activity compared to tissues from control patients, as demonstrated by phospho-p65 staining (p<0.0001; Figure 6E-F). Together, these data suggest that epithelial hyperplasia in BPH may be promoted by inflammation-derived TNFα and that this can be abrogated by systemic treatment with TNFα-antagonists.

**Figure 6.**
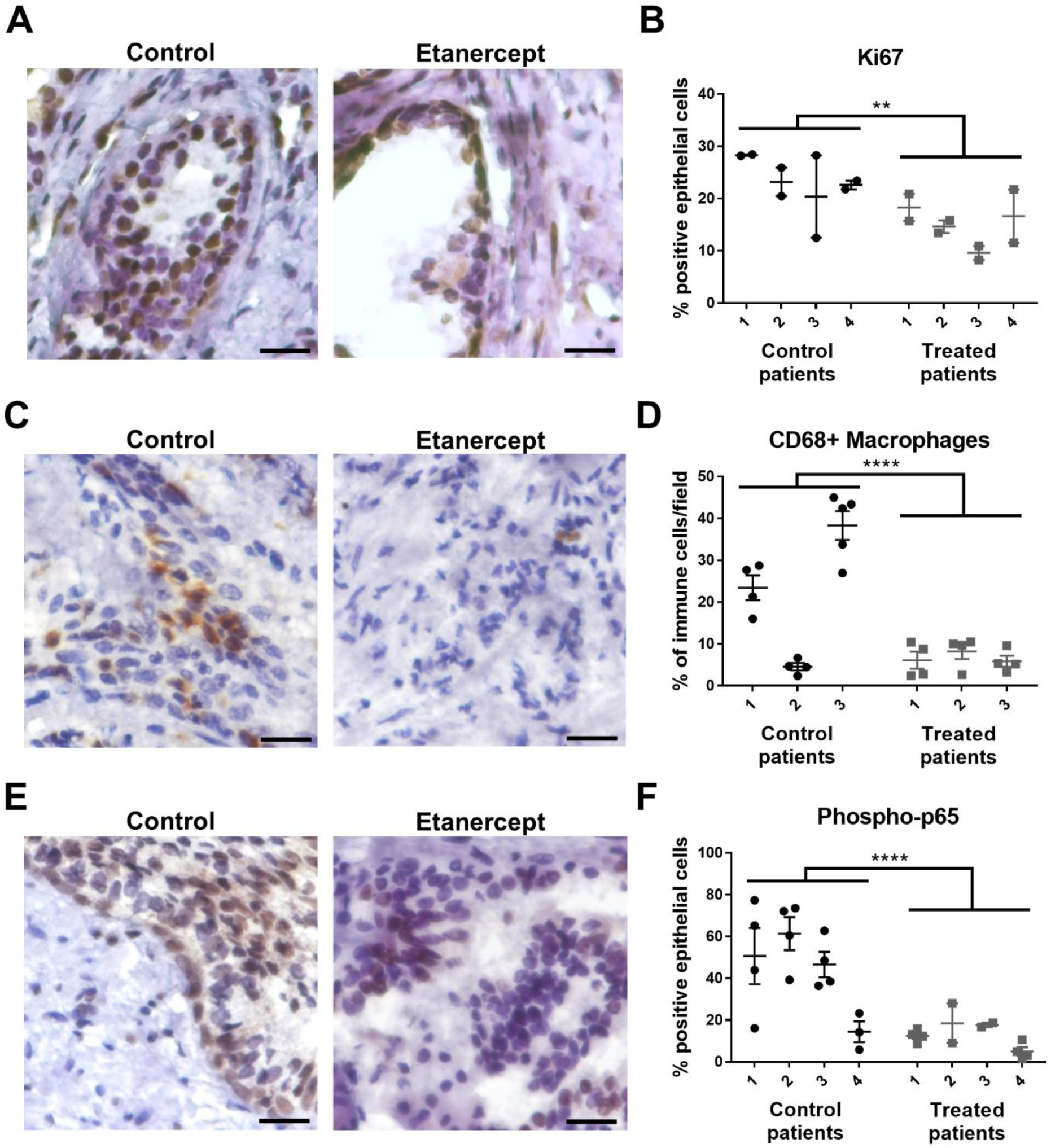
Patients treated with TNFα-antagonists have decreased prostatic epithelial proliferation, macrophage infiltration, and NFκB activation. Human transition zone tissues from patients taking TNFα-antagonists at the time of radical prostatectomy or matched controls were used for histological evaluation. **A)** Representative images of IHC staining for Ki67 in control or treated patients. **B)** Quantitation of IHC staining for epithelial Ki67 in control or treated patients, indicated as the percent Ki67+ epithelial cells per field. **C)** Representative images of IHC staining for CD68 in control or treated patients. **D)** IHC quantitation indicating the abundance of CD68+ macrophages, represented as the portion of all immune cells per field. **E)** Representative images of phospho-p65 staining via IHC in control or treated patients. **F)** Data represents the quantitation of IHC staining for epithelial phospho-p65 staining in control or treated patients, represented as the percent positive epithelial cells per field. **B, D, F)** Data indicate the mean +/- SEM of the percent positive cells per field, where individual points indicate separate fields for each patient. Comparison of control and TNFα-antagonist treated groups were conducted using a mixed effects model asterisks indicate the significance of the treatment. Scale bars = 20µm.

## Discussion

The present results suggest a new biological indication for the use of TNFα-antagonists, a well-tolerated class of drugs commonly used to treat AI disease, as a mechanism to alter the pathogenesis of BPH. These agents, but not methotrexate, reduce BPH incidence in patients with AI diseases. In animal models, TNFα-antagonists reduce both the establishment of epithelial hyperplasia and, in aged mice, prostate size. A number of trials using NSAIDs to relieve BPH symptoms have shown some positive effects, but limited long-term efficacy (*42, 43*). The present data strongly suggest that appropriate targeting of specific molecular signals may be more clinically effective than broad-spectrum agents. There are many immunomodulatory drugs in development or already being prescribed for AI diseases that could be tested for their activity in BPH.

These studies strengthen the link between prostate inflammation and the development of BPH. NFκB activation has been shown to be increased in BPH samples progressing to disease-specific surgery and was associated with increased expression of 5α-reductase 2, increased androgen receptor expression, and increased cellular proliferation *in vitro* (*44, 45*), and TNFα may drive this paradigm. Inflammation can directly affect molecular pathways associated with growth and resistance to 5ARI therapies (*44, 45*). Furthermore, natural remedies such as saw palmetto and beta-sitosterol have also been suggested to reduce NFκB activity (*46, 47*), indicating that NFκB activation may be involved in driving and/or supporting hyperplastic growth.

The immunosuppressive function of androgens has been reported in numerous studies (*32*). The contributing role(s) of the androgen receptor in BPH is not fully understood, and the involvement of androgen receptor signaling in prostatic inflammation may contribute to this complexity (*48, 49*). Reduced circulating androgen levels in aging men might contribute to increased inflammation in BPH, but whether the increase in inflammation is in response to a stimulus (e.g. wound repair) or is autoimmune-mediated remains to be determined. It is also possible that TNFα secreted by inflammatory cells alters prostatic androgen/estrogen levels since this cytokine is known to stimulate aromatase activity; TNFα-antagonist treatment in RA decreases aromatase action and increases synovial androgen levels (*50*).

Since BPH patients have one or more of a variety of histological features, including stromal nodules, glandular nodules, and/or fibrosis, this disease is more likely a combination of conditions (*51*). Recent molecular profiling studies have provided a basis for at least two BPH subtypes (*12*). As in prostate development, it has long been clear that paracrine interactions between stromal and epithelial tissues are important in BPH (*41*). Appreciating an additional role for immune cells in this intercellular conversation increases the perceived complexity of these interactions but opens new therapeutic targets and will be critical to elucidating mechanisms by which TNFα or other signaling pathways drive hyperplasia and/or fibrosis. Whether inflammation drives a specific subtype of BPH is not clear, but it is certainly possible that inflammatory factors influence many features of BPH, including hyperplasia and collagen deposition (*52–54*).

The use of scRNA-seq analyses in both characterizing the cell types present in and providing a molecular understanding of diseases have been extremely useful in moving biological studies forward. Evaluation of cell populations from synovial tissue in RA patients indicate that while different cell types may express either *TNF* or *TNFRSF1A* (TNFR1), only monocytes expressed high levels of both of these genes (*33*). Furthermore, inflammatory monocytes in ulcerative colitis and Crohn’s disease express *TNF* but may also aid in anti-TNFα therapy resistance (*34, 35*). Even though it is clear that TNFα signaling is important in AI conditions and BPH, numerous cell types and/or inflammatory pathways can be explored for therapeutic potential using these scRNA-seq datasets. Each AI disease is distinct, with different immune cell activation states, stromal cell populations, and levels of systemic inflammation. Whether the inflammatory cell populations in BPH closely mirror those of an individual AI disease remains to be determined (*33–35, 55*).

Due to the retrospective nature of these studies, there are several limitations. While this population represents a large cohort of men with BPH, the subset of patients undergoing treatment for AI disease with a concomitant diagnosis of BPH and/or have progressed to surgery for prostatic disease was quite small. The samples used were dependent on the surgical procedure and the prostate biorepository has tissues from less than 800 patients. Expanding the patient population will determine the capacity for TNFα-antagonists to reduce surgical endpoints for BPH as well as discern if there are associations with AI disease treatment and BPH diagnoses across racial and ethnic groups. Another limitation of these studies is that the specific cell types that contribute to BPH through direct *versus* indirect responses to TNFα have yet to be determined *in vivo*. Obvious limitations using *in vitro* assays, including a lack of complexity compared to the *in vivo* BPH microenvironment, warrant further investigation of TNFα function in the prostate. Additionally, no true murine model for BPH exists, so the NOD and Pb-PRL mouse models used for these studies lack some aspects of human disease (such as fibrosis and the influence of androgen:estrogen ratios) and may limit the translational potential of future mechanistic studies. Finally, since the occurrence of a prostatectomy in men taking TNFα-antagonists is rare, so scRNA-seq studies to-date have not been able to directly evaluate the impact of these drugs on the prostate’s inflammatory milieu and proliferative prostatic cell types.

It is likely these drugs would be most beneficial to patients with chronic prostatic inflammation, so it would also be useful to detect intraprostatic inflammation through non-invasive imaging procedures (*56, 57*) or to be able to determine the risk of development of BPH through personalized medicine and genetic risk scores (*11*). Further evaluation of human BPH tissues will determine if the presence of inflammatory macrophages or other identified cell signatures could contribute to anti-TNFα therapy resistance (*35*). The effect of TNFα in stromal versus glandular BPH nodules also remains to be determined. Importantly, these antagonists are already widely used and future work will determine whether short-term treatments (to limit the side effects of systemic anti-inflammatory agents) or potentially combination with current BPH therapies are beneficial. Such studies will provide a basis for additional urological evaluation via earlier monitoring and therapeutic intervention in men with AI diseases. Tailoring the AI disease treatment strategy to include limiting BPH incidence could add therapeutic benefit for men. Such a strategy could both enhance patient quality of life and decrease the need for BPH surgical endpoints in these patients.

## Materials and Methods

The study design for each experiment is included within respective subsections.

### Enterprise Database Warehouse Human Subjects Study

A retrospective evaluation of patients was conducted for 112,152 males with office visits at NorthShore University HealthSystem between 01/01/2010 and 12/31/2012. A preliminary analysis determined that an expanded dataset should be used for final analysis. Patients under 40 years of age or with a diagnosis of prostate cancer were excluded from the study. Records were searched for patient diagnoses of BPH or any AI conditions (Table S1). Baseline BPH incidence rates were identified with a Chi-square test. Chi-square tests were utilized to compare the proportion of men with BPH by autoimmune condition to the proportion of men with BPH and no AI condition, as well as to compare the proportion of BPH diagnoses in men being treated for an AI condition to the proportion of men with BPH and no AI condition. Predictors of BPH diagnosis were tested using multivariable logistic regression models and predictors were determined a priori. For men who underwent surgery for BPH, median days to surgery by medication use were compared using the Mann-Whitney U test. Statistical significance was established throughout at p<0.05. Statistical analyses were conducted using SAS version 9.4 (SAS Institute Inc, Cary, NC).

### Isolation of CD45+ cells from human tissues

Human prostate tissues were procured with the IRB-approved NorthShore Urologic Disease Biorepository and Database with informed consent and de-identified clinical annotation. Small prostate tissues were obtained from patients undergoing robotic-assisted laparoscopic prostatectomy (RALP) for prostate cancer, International Prostate Symptom Score (IPSS) of <15, Gleason 6-7, and estimated prostate volume of <40 grams by imaging with transrectal ultrasound (TRUS), CT scan, or MRI. Large prostate tissues were obtained from patients undergoing either RALP for prostate cancer (Gleason 6-7) or simple prostatectomy for BPH and had an estimated prostate size of >90 grams. The transition zone (TZ) was dissected and separated for formalin-fixed paraffin-embedded (FFPE) histology or digested and prepared for fluorescence activated cell sorting (FACS). Tissues were minced, then digested while shaking at 37°C for 2 hours in 200 U/mL Collagenase I (Gibco) + 1 mg/mL DNase I (Roche) + 1% antibiotic/antimycotic solution in Hank’s Balanced Salt Solution. Digestion solution was replaced with TrypLE express dissociation reagent (Gibco) and allowed to shake at 37°C for 5-10 minutes. Digested samples were neutralized in RPMI + 10% FBS, then mechanically disrupted by pipetting repeatedly. Samples were passed through a 100µm cell strainer, then washed. Red blood cells were lysed in a hypotonic buffer, then cells were stained with Zombie Violet (Biolegend) and blocked with Human TruStain FcX blocking antibody (Biolegend). CD45-PE [clone HI30], EpCAM-APC [clone 9C4], and CD200- PE/Cy7 [clone OX-104] antibodies (Biolegend) were added to stain samples in preparation for FACS on a BD FACSAria II. Approximately 100,000 viable CD45+CD200-EpCAM- cells were sorted for downstream analysis.

### scRNA-seq of CD45+ cells

FACS-isolated cells were spun down and washed at least twice prior to loading onto the 10X Chromium System (10X Genomics), with Single Cell 3’ Library & Gel Bead Kit, v3.0 reagents. Cells from three small and three large tissues were stained with TotalSeq-B Antibodies (Biolegend) for CITE-seq analysis. Antibodies for CD3 [clone UCHT1], CD4 [clone RPA-T4], CD8 [clone RPA-T8], CD19 [clone HIB19], and CD11b [clone ICRF44] were used following the manufacturer’s instructions prior to loading into the Chromium System. Cells were loaded for downstream evaluation of 5,000 cells/sample and cDNA amplification and library preparation were conducted according to the manufacturer’s instructions. Libraries were sent to the Purdue Genomics Core Facility for post-library construction quality control, quantification, and sequencing. A high sensitivity DNA chip was run on an Agilent Bioanalyzer (Agilent) per the recommendation of 10x Genomics. Additional quality control was performed by running a denatured DNA pico chip (Agilent) followed by an AMPure cleanup (Beckman Coulter). Final library quantification was completed using a Kapa kit (KAPA Biosystems) prior to sequencing. Sequencing was conducted using a NovaSeq S4 flow cell on a NovaSeq 6000 system (Illumina) with 2x150 base-pair reads at a depth of 50,000 reads/cell. Libraries generated from cell surface protein labeling with TotalSeq-B antibodies were sequenced at a depth of 5,000 reads/cell.

### Data processing and quality control

Sequencing reads from the Chromium system were de-multiplexed and processed using the CellRanger pipeline v3.0.0 (10x Genomics). CellRanger mkfastq was run to generate FASTQ files where dual indices were ignored, barcode mismatch allowance was set to 0, and the flag “— use-bases-mask=Y26n*,I8n*,n*,Y98n” was set. CellRanger count was then used for alignment, filtering, barcode counting, and unique molecular identifier (UMI) counting. All reads were aligned to the ENSEMBL human genome version GrCh38 using the STAR aligner v2.5.4 (*58*). CellRanger was run with the expected cells set to 5,000.

R version 3.5.1 and Bioconductor version 3.8 were used in all statistical analyses. Cells that had fewer than 1,000 or greater than 10,000 observed genes were discarded. Cells were also removed if greater than 22% of all reads mapped to mitochondrial genes. A summary of the data produced by the scRNA-seq analyses, including the run metrics, is shown in Table S4.

### Unsupervised clustering and identification of marker genes

All scRNA-seq data were uploaded to GEO and are available with accession number GSE164695. Seurat version 3.1.3 was used for data normalization and cell clustering based on differential gene expression (*59, 60*). Data were normalized using scTransform (*61*) v.0.3.1 and cell cycle-related genes were used to produce a cell cycle score for each cell. Cell cycle scores, mitochondrial reads, and UMI counts were used to regress out heterogeneity from these variables by scaling the data. These “corrected” data, after permutation and selection of the first 30 principal components based on principal component analysis (PCA) scores, were used for downstream analysis. Unsupervised clustering was performed in Seurat, which uses graph-based approaches to first construct K-nearest neighbor graphs (K = 30) and identifies clusters by iteratively forming communities of cells to optimize the modularity function. The number of clusters were determined using the Louvain method (*62*) for community detection, as implemented in Seurat with a resolution of 0.2. The correct resolution to use was determined both visually through plots and heat maps as well as using clustering trees via the clustree (*63*) R package v 0.4.3, selecting a resolution that provides stable clusters. P-values were corrected for multiple testing using the Benjamini-Hochberg method (*64*). Biomarkers were considered statistically significant at a 1% false discovery rate (FDR) using the Wilcoxon rank sum test (*65*). Differentially expressed genes between small and large sample groups were identified using the edgeR (*66, 67*) Bioconductor package, v 3.31 with an FDR cutoff of 5%.

### Animal Studies

Male non-obese diabetic (NOD) mice were purchased from Jackson Laboratory (Bar Harbor, ME; Stock Number: 001976) and maintained in a barrier animal facility at NorthShore University HealthSystem. All studies were conducted according to US federal and state regulations and approved by the NorthShore Institutional Animal Care and use Committee (IACUC). Mice age 25 weeks were injected with 4 mg/kg TNFα antagonist etanercept (Enbrel®; n=12) or vehicle (PBS, n=8) twice weekly by intra-peritoneal injection for five weeks. Mice were randomly assigned to groups using block stratification. Diabetic status was tested using a glucometer, as previously described (*23*), at the time of first injection and last injection. Mice were harvested at 30 weeks.

Pb-PRL transgenic mice were from Dr. Kindblom, Sahlgrenska University Hospital. All studies were performed in accordance with the National Institute of Health Guidelines for the Care and Use of Laboratory Animals and approved by the Roswell Park Institutional Animal Care and Use Committee (#1308M). Male Pb-PRL mice (20-22 months) were treated with 4 mg/kg TNF ligand trap, etanercept (Enbrel®, n = 5), or with vehicle (PBS, n = 6) twice a week by intra- peritoneal injection for 12 weeks. The ventral prostate volume was measured every four weeks using high-resolution, high-frequency ultrasound, as described in (*40*). Data is presented as mean +/- SEM, normalized to pre-treatment volume (151 +/-10 mm^3^). One animal in the PBS group was removed from the analysis for week 12 due to rapid prostate swelling, determined on necropsy to likely have arisen from a hemorrhage. Thus, data indicate n=5 for each group at week 12. In both animal models, urogenital tract tissues were harvested and formalin-fixed, paraffin-embedded (FFPE) in preparation for histology.

### Immunohistochemistry

Tissue sections of 5µm thickness were mounted on slides and prepared for immunohistochemistry (IHC) staining as previously described (*23*). Detailed information on antibodies used for staining in human and mouse tissues is included in Table S5. Quantitation of Ki67+ or phospho-p65+ cells was performed within the epithelial cell compartment and F4/80+ (mouse) or CD68+ (human) macrophages were evaluated within the immune cell compartment. All human and mouse IHC quantitation was completed by S.C. by counting the indicated number of fields under the 40x objective for a random subset of samples and included tissues from three or more patients/animals per stain. In figures, the sample numbers in one panel do not necessarily correspond to the numbers in another panel.

### Cell Culture

THP-1 cells were purchased and authenticated from ATCC (STRB0424) and used within 20 passages of acquisition. BHPrE-1, NHPrE-1, and BHPrS-1 cell lines were isolated and cultured as benign epithelial and stromal prostatic cell models, as described (*68, 69*). Authentication of BHPrE-1 (STRA3426), NHPrE-1 (STRA3441), and BHPrS-1 (STRB0418) cells was completed by ATCC and all experiments were conducted within 20 passages of testing.

THP-1 cells were cultured exactly as indicated by ATCC and were polarized as described previously (*70*). Briefly, THP-1 cells were differentiated for 24 hours with PMA, followed by M1 polarization with LPS + IFNγ for 24 hours or M2 polarization with IL-4 + IL-13 for 72 hours. After polarization was complete, serum free medium was added to M1 or M2 macrophages and incubated for 24 hours. Conditioned medium was then harvested, filtered, and stored at -80°C until it was used for growth assays.

Isolation of primary fibroblasts from de-identified BPH patient tissues 012 and 376 was completed as previously described (*71*). Briefly, freshly isolated human prostate transition zone tissue was obtained after simple prostatectomy followed by mechanical and enzymatic digestion. Fibroblasts were cultured and used for all assays prior to passage 12.

For crystal violet growth assays, cells were allowed to adhere to 96-well plates prior to indicated treatments. Human recombinant TNFα was purchased from Peprotech and TNFα- neutralizing antibody (P300A) was pre-incubated at 40 µg/mL with 50% macrophage conditioned medium for 4-6 hours. Cells were grown in low serum (0.1%) with treatment conditions for up to six days and cells fixed with 4% paraformaldehyde. Cells were stained with crystal violet and washed, then stain solubilized in 10% acetic acid to obtain relative absorbance values.

### Statistical Analysis

Statistical significance of *in vitro* assays was completed using a two-way analysis of variance (ANOVA), patient characteristics were compared using a student’s *t*-test, and immune cell compartments were correlated using linear regression using Prism software version 7.05 (GraphPad). A mixed effects model was used for IHC quantitation evaluation using SAS 9.4 (Cary, NC). A p-value of less than 0.05 was considered significant. In data figures, significance is indicated by *=p<0.05, **=p<0.01, ***=p<0.001, and ***=p<0.0001.

## Supplementary Materials

**Figure S1.** Fluorescence activated cell sorting strategy for CD45+ scRNA-seq analysis.

**Figure S2.** Additional data relating to scRNA-seq data and patient characteristics.

**Figure S3.** Estimated proportions of immune cell types by scRNA-seq analysis represent immune cell proportions by flow cytometry.

**Figure S4.** Expression level of TNF, TNFRSF1A, and TNFRSF1B in BPH leukocytes.

**Figure S5.** TNFα-antagonist treatment in Pb-PRL mice does not significantly alter macrophage infiltration or epithelial NFκB activity.

**Figure S6.** TNF promotes stromal, but not epithelial, cell growth.

**Figure S7.** Characteristics of TNFα-antagonist treated or control patients.

**Table S1.** Proportion of men with BPH by autoimmune disease status.

**Table S2.** Median Days for Progression to Surgery for BPH Patients.

**Table S3.** Most significantly altered pathways in CD45+ cells from large versus small human BPH tissues.

**Table S4.** Run metrics for scRNA-seq of BPH associated CD45+ leukocytes. Table S5. Antibodies used for IHC in human and mouse tissues.

## Acknowledgements

The authors appreciate the assistance of Catherine Zhu in extracting medical record data and the Purdue University Genomics Facility for their aid in scRNA-seq library normalization and sequencing. The authors are grateful to the NorthShore Biospecimen Repository and patients who have donated their tissue for research, without which much of this work would not have been possible. **Funding:** This work was funded by 1P20DK116185 (S.W.H. and T.L.R.) and R01DK117906 (S.W.H.) from NIDDK, the Purdue University Center for Cancer Research (NIH grant P30CA023168), the IU Simon Cancer Center (NIH grant P30CA082709), and the Roswell Park Comprehensive Cancer Center and imaging facility (NIH grants P30CA016056 and S10OD010393-01). This work was also generously supported by the Collaborative Core for Cancer Bioinformatics, the Walther Cancer Foundation, and the Rob Brooks Fund for Precision Prostate Cancer Care. **Author Contributions:** S.E.C., O.E.F., and S.W.H. created the study concept; R.E.V., S.E.C., O.E.F., T.L.R., K.L.N. and S.W.H. designed research studies; R.E.V., J.P., P.T., C.Z., B.T.H., A.P.G., O.E.F., and S.W.H. were involved in acquisition of patient tissues or accessing medical records; B.L., C-H.W., and N.A.L. performed statistical and bioinformatics analyses; R.E.V. L.A-B., R.Z., B.L., V.G., M.G., O.E.F., G.M.C., M.M.B., T.S., and S.E.C. performed data acquisition and all authors contributed to data analysis and interpretation. R.E.V., S.E.C., and S.W.H. wrote the manuscript and all authors critically revised the manuscript. **Competing interests:** None of the authors have competing interests to disclose. **Data and materials availability:** The scRNA-seq data are available in the NCBI GEO repository (accession GSE164695).

## References

1. S. J. Berry, D. S. Coffey, P. C. Walsh, and L. L. Ewing. The development of human benign prostatic hyperplasia with age. J Urol 132, 474–479 (1984).

2. K. T. McVary. BPH: epidemiology and comorbidities. Am J Manag Care 12, S122–128 (2006).

3. T. M. Nicholson, and W. A. Ricke. Androgens and estrogens in benign prostatic hyperplasia: past, present and future. Differentiation 82, 184–199 (2011).

4. J. H. Fowke, T. Koyama, O. Fadare, and P. E. Clark. Does Inflammation Mediate the Obesity and BPH Relationship? An Epidemiologic Analysis of Body Composition and Inflammatory Markers in Blood, Urine, and Prostate Tissue, and the Relationship with Prostate Enlargement and Lower Urinary Tract Symptoms. PLoS One 11, e0156918 (2016).

5. H. Nandeesha, B. C. Koner, L. N. Dorairajan, and S. K. Sen. Hyperinsulinemia and dyslipidemia in non-diabetic benign prostatic hyperplasia. Clin Chim Acta 370, 89–93 (2006).

6. J. K. Parsons, A. V. Sarma, K. McVary, and J. T. Wei. Obesity and benign prostatic hyperplasia: clinical connections, emerging etiological paradigms and future directions. J Urol 189, S102–106 (2013).

7. A. V. Sarma, J. L. St Sauver, J. M. Hollingsworth, D. J. Jacobson, M. E. McGree, R. L. Dunn, M. M. Lieber, S. J. Jacobsen, and P. Urologic Diseases in America. Diabetes treatment and progression of benign prostatic hyperplasia in community-dwelling black and white men. Urology 79, 102–108 (2012).

8. S. K. Van Den Eeden, A. Ferrara, J. Shan, S. J. Jacobsen, V. P. Quinn, R. Haque, and C. P. Quesenberry. Impact of type 2 diabetes on lower urinary tract symptoms in men: a cohort study. BMC Urol 13, 12 (2013).

9. K. S. Coyne, S. A. Kaplan, C. R. Chapple, C. C. Sexton, Z. S. Kopp, E. N. Bush, L. P. Aiyer, and L. T. Epi. Risk factors and comorbid conditions associated with lower urinary tract symptoms: EpiLUTS. BJU Int 103 **Suppl 3**, 24–32 (2009).

10. J. N. Hellwege, S. Stallings, E. S. Torstenson, R. Carroll, K. M. Borthwick, M. H. Brilliant, D. Crosslin, A. Gordon, G. Hripcsak, G. P. Jarvik, J. G. Linneman, P. Devi, P. L. Peissig, P. A. M. Sleiman, H. Hakonarson, M. D. Ritchie, S. S. Verma, N. Shang, J. C. Denny, D. M. Roden, D. R. Velez Edwards, and T. L. Edwards. Heritability and genome-wide association study of benign prostatic hyperplasia (BPH) in the eMERGE network. Sci Rep 9, 6077 (2019).

11. . A. J. Gudmundsson, J. K. Sigurdsson, L. Stefansdottir, B. A. Agnarsson, H. J. Isaksson, O. Stefansson, S. A. Gudjonsson, D. F. Gudbjartsson, G. Masson, M. L. Frigge, S. N. Stacey, P. Sulem, G. H. Halldorsson, V. Tragante, H. Holm, G. I. Eyjolfsson, O. Sigurdardottir, I. Olafsson, T. Jonsson, E. Jonsson, R. B. Barkardottir, R. Hilmarsson, F. W. Asselbergs, G. Geirsson, U. Thorsteinsdottir, T. Rafnar, G. Thorleifsson, and K. Stefansson. Genome-wide associations for benign prostatic hyperplasia reveal a genetic correlation with serum levels of PSA. Nat Commun 9, 4568 (2018).

12. D. Liu, J. E. Shoag, D. Poliak, R. S. Goueli, V. Ravikumar, D. Redmond, A. Vosoughi, J. Fontugne, H. Pan, D. Lee, D. Thomas, K. Salari, Z. Wang, A. Romanel, A. Te, R. Lee, B. Chughtai, A. F. Olumi, J. M. Mosquera, F. Demichelis, O. Elemento, M. A. Rubin, A. Sboner, and C. E. Barbieri. Integrative multiplatform molecular profiling of benign prostatic hyperplasia identifies distinct subtypes. Nat Commun 11, 1987 (2020).

13. J. D. McConnell, C. G. Roehrborn, O. M. Bautista, G. L. Andriole, Jr., C. M. Dixon, J. W. Kusek, H. Lepor, K. T. McVary, L. M. Nyberg, Jr., H. S. Clarke, E. D. Crawford, A. Diokno, J. P. Foley, H. E. Foster, S. C. Jacobs, S. A. Kaplan, K. J. Kreder, M. M. Lieber, M. S. Lucia, G. J. Miller, M. Menon, D. F. Milam, J. W. Ramsdell, N. S. Schenkman, K. M. Slawin, and J. A. Smith. The long-term effect of doxazosin, finasteride, and combination therapy on the clinical progression of benign prostatic hyperplasia. N Engl J Med 349, 2387–2398 (2003).

14. L. Feinstein, and B. Matlaga. Urologic Diseases in America. **NIH Publication** No. 12- 7865, 8–37 (2018).

15. K. T. McVary. A review of combination therapy in patients with benign prostatic hyperplasia. Clin Ther 29, 387–398 (2007).

16. J. E. Elkahwaji, W. Zhong, W. J. Hopkins, and W. Bushman. Chronic bacterial infection and inflammation incite reactive hyperplasia in a mouse model of chronic prostatitis. Prostate 67, 14–21 (2007).

17. W. A. Bushman, and T. J. Jerde. The role of prostate inflammation and fibrosis in lower urinary tract symptoms. Am J Physiol Renal Physiol 311, F817–F821 (2016).

18. D. W. Strand, L. Aaron, G. Henry, O. E. Franco, and S. W. Hayward. Isolation and analysis of discreet human prostate cellular populations. Differentiation 91, 139–151 (2016).

19. X. Wang, W. J. Lin, K. Izumi, Q. Jiang, K. P. Lai, D. Xu, L. Y. Fang, T. Lu, L. Li, S. Xia, and C. Chang. Increased infiltrated macrophages in benign prostatic hyperplasia (BPH): role of stromal androgen receptor in macrophage-induced prostate stromal cell proliferation. J Biol Chem 287, 18376–18385 (2012).

20. G. Kramer, D. Mitteregger, and M. Marberger. Is benign prostatic hyperplasia (BPH) an immune inflammatory disease? Eur Urol 51, 1202–1216 (2007).

21. Y. M. Tzeng, L. T. Kao, H. C. Lin, and C. Y. Huang. A Population-Based Study on the Association between Benign Prostatic Enlargement and Rheumatoid Arthritis. PLoS ONE 10, e0133013 (2015).

22. O. Lin-Tsai, P. E. Clark, N. L. Miller, J. H. Fowke, O. Hameed, S. W. Hayward, and D. W. Strand. Surgical intervention for symptomatic benign prostatic hyperplasia is correlated with expression of the AP-1 transcription factor network. Prostate 74, 669–679 (2014).

23. L. M. Aaron-Brooks, T. Sasaki, R. E. Vickman, L. Wei, O. E. Franco, Y. Ji, S. E. Crawford, and S. W. Hayward. Hyperglycemia and T Cell infiltration are associated with stromal and epithelial prostatic hyperplasia in the nonobese diabetic mouse. Prostate 79, 980–993 (2019).

24. E. Gremese, B. Tolusso, M. R. Gigante, and G. Ferraccioli. Obesity as a risk and severity factor in rheumatic diseases (autoimmune chronic inflammatory diseases). Front Immunol 5, 576 (2014).

25. J. Zhang, L. Fu, J. Shi, X. Chen, Y. Li, B. Ma, and Y. Zhang. The risk of metabolic syndrome in patients with rheumatoid arthritis: a meta-analysis of observational studies. PLoS One 8, e78151 (2013).

26. A. Dregan, J. Charlton, P. Chowienczyk, and M. C. Gulliford. Chronic inflammatory disorders and risk of type 2 diabetes mellitus, coronary heart disease, and stroke: a population-based cohort study. Circulation 130, 837–844 (2014).

27. D. Furman, J. Campisi, E. Verdin, P. Carrera-Bastos, S. Targ, C. Franceschi, L. Ferrucci, D. W. Gilroy, A. Fasano, G. W. Miller, A. H. Miller, A. Mantovani, C. M. Weyand, N. Barzilai, J. J. Goronzy, T. A. Rando, R. B. Effros, A. Lucia, N. Kleinstreuer, and G. M. Slavich. Chronic inflammation in the etiology of disease across the life span. Nat Med 25, 1822–1832 (2019).

28. J. J. Wu, T. U. Nguyen, K. Y. Poon, and L. J. Herrinton. The association of psoriasis with autoimmune diseases. J Am Acad Dermatol 67, 924–930 (2012).

29. M. Feldmann, and R. N. Maini. Lasker Clinical Medical Research Award. TNF defined as a therapeutic target for rheumatoid arthritis and other autoimmune diseases. Nat Med 9, 1245–1250 (2003).

30. N. Gleicher, and D. H. Barad. Gender as risk factor for autoimmune diseases. J Autoimmun 28, 1–6 (2007).

31. M. Benagiano, P. Bianchi, M. M. D’Elios, I. Brosens, and G. Benagiano. Autoimmune diseases: Role of steroid hormones. Best Pract Res Clin Obstet Gynaecol 60, 24–34 (2019).

32. V. E. Bianchi. The Anti-Inflammatory Effects of Testosterone. J Endocr Soc 3, 91–107 (2019).

33. F. Zhang, K. Wei, K. Slowikowski, C. Y. Fonseka, D. A. Rao, S. Kelly, S. M. Goodman, D. Tabechian, L. B. Hughes, K. Salomon-Escoto, G. F. M. Watts, A. H. Jonsson, J. Rangel-Moreno, N. Meednu, C. Rozo, W. Apruzzese, T. M. Eisenhaure, D. J. Lieb, D. L. Boyle, A. M. Mandelin, 2nd, A. Accelerating Medicines Partnership Rheumatoid, C. Systemic Lupus Erythematosus, B. F. Boyce, E. DiCarlo, E. M. Gravallese, P. K. Gregersen, L. Moreland, G. S. Firestein, N. Hacohen, C. Nusbaum, J. A. Lederer, H. Perlman, C. Pitzalis, A. Filer, V. M. Holers, V. P. Bykerk, L. T. Donlin, J. H. Anolik, M. B. Brenner, and S. Raychaudhuri. Defining inflammatory cell states in rheumatoid arthritis joint synovial tissues by integrating single-cell transcriptomics and mass cytometry. Nat Immunol 20, 928–942 (2019).

34. C. S. Smillie, M. Biton, J. Ordovas-Montanes, K. M. Sullivan, G. Burgin, D. B. Graham, R. H. Herbst, N. Rogel, M. Slyper, J. Waldman, M. Sud, E. Andrews, G. Velonias, A. L. Haber, K. Jagadeesh, S. Vickovic, J. Yao, C. Stevens, D. Dionne, L. T. Nguyen, A. C. Villani, M. Hofree, E. A. Creasey, H. Huang, O. Rozenblatt-Rosen, J. J. Garber, H. Khalili, A. N. Desch, M. J. Daly, A. N. Ananthakrishnan, A. K. Shalek, R. J. Xavier, and A. Regev. Intra- and Inter-cellular Rewiring of the Human Colon during Ulcerative Colitis. Cell 178, 714–730 e722 (2019).

35. J. C. Martin, C. Chang, G. Boschetti, R. Ungaro, M. Giri, J. A. Grout, K. Gettler, L. S. Chuang, S. Nayar, A. J. Greenstein, M. Dubinsky, L. Walker, A. Leader, J. S. Fine, C. E. Whitehurst, M. L. Mbow, S. Kugathasan, L. A. Denson, J. S. Hyams, J. R. Friedman, P. T. Desai, H. M. Ko, I. Laface, G. Akturk, E. E. Schadt, H. Salmon, S. Gnjatic, A. H. Rahman, M. Merad, J. H. Cho, and E. Kenigsberg. Single-Cell Analysis of Crohn’s Disease Lesions Identifies a Pathogenic Cellular Module Associated with Resistance to Anti-TNF Therapy. Cell 178, 1493–1508 e1420 (2019).

36. M. Stoeckius, C. Hafemeister, W. Stephenson, B. Houck-Loomis, P. K. Chattopadhyay, H. Swerdlow, R. Satija, and P. Smibert. Simultaneous epitope and transcriptome measurement in single cells. Nat Methods 14, 865–868 (2017).

37. H. Wennbo, J. Kindblom, O. G. Isaksson, and J. Tornell. Transgenic mice overexpressing the prolactin gene develop dramatic enlargement of the prostate gland. Endocrinology 138, 4410–4415 (1997).

38. J. Kindblom, K. Dillner, L. Sahlin, F. Robertson, C. Ormandy, J. Tornell, and H. Wennbo. Prostate hyperplasia in a transgenic mouse with prostate-specific expression of prolactin. Endocrinology 144, 2269–2278 (2003).

39. J. S. Davis, K. L. Nastiuk, and J. J. Krolewski. TNF is necessary for castration-induced prostate regression, whereas TRAIL and FasL are dispensable. Mol Endocrinol 25, 611–620 (2011).

40. S. Singh, C. Pan, R. Wood, C. R. Yeh, S. Yeh, K. Sha, J. J. Krolewski, and K. L. Nastiuk. Quantitative volumetric imaging of normal, neoplastic and hyperplastic mouse prostate using ultrasound. BMC Urol 15, 97 (2015).

41. J. E. McNeal. Anatomy of the prostate and morphogenesis of BPH. Prog Clin Biol Res 145, 27–53 (1984).

42. J. M. Schenk, G. S. Calip, C. M. Tangen, P. Goodman, J. K. Parsons, I. M. Thompson, and A. R. Kristal. Indications for and use of nonsteroidal antiinflammatory drugs and the risk of incident, symptomatic benign prostatic hyperplasia: results from the prostate cancer prevention trial. Am J Epidemiol 176, 156–163 (2012).

43. J. L. St Sauver, D. J. Jacobson, M. E. McGree, M. M. Lieber, and S. J. Jacobsen. Protective association between nonsteroidal antiinflammatory drug use and measures of benign prostatic hyperplasia. Am J Epidemiol 164, 760–768 (2006).

44. D. C. Austin, D. W. Strand, H. L. Love, O. E. Franco, M. M. Grabowska, N. L. Miller, O. Hameed, P. E. Clark, R. J. Matusik, R. J. Jin, and S. W. Hayward. NF-kappaB and androgen receptor variant 7 induce expression of SRD5A isoforms and confer 5ARI resistance. Prostate 76, 1004–1018 (2016).

45. D. C. Austin, D. W. Strand, H. L. Love, O. E. Franco, A. Jang, M. M. Grabowska, N. L. Miller, O. Hameed, P. E. Clark, J. H. Fowke, R. J. Matusik, R. J. Jin, and S. W. Hayward. NF-kappaB and androgen receptor variant expression correlate with human BPH progression. Prostate 76, 491–511 (2016).

46. H. V. Sudeep, K. Venkatakrishna, B. Amrutharaj, Anitha, and K. Shyamprasad. A phytosterol-enriched saw palmetto supercritical CO2 extract ameliorates testosterone-induced benign prostatic hyperplasia by regulating the inflammatory and apoptotic proteins in a rat model. BMC Complement Altern Med 19, 270 (2019).

47. Y. Sun, L. Gao, W. Hou, and J. Wu. beta-Sitosterol Alleviates Inflammatory Response via Inhibiting the Activation of ERK/p38 and NF-kappaB Pathways in LPS-Exposed BV2 Cells. Biomed Res Int 2020, 7532306 (2020).

48. R. E. Vickman, O. E. Franco, D. C. Moline, D. J. Vander Griend, P. Thumbikat, and S. W. Hayward. The role of the androgen receptor in prostate development and benign prostatic hyperplasia: A review. Asian J Urol 7, 191–202 (2020).

49. B. Cioni, A. Zaalberg, J. R. van Beijnum, M. H. M. Melis, J. van Burgsteden, M. J. Muraro, E. Hooijberg, D. Peters, I. Hofland, Y. Lubeck, J. de Jong, J. Sanders, J. Vivie, H. G. van der Poel, J. P. de Boer, A. W. Griffioen, W. Zwart, and A. M. Bergman. Androgen receptor signalling in macrophages promotes TREM-1-mediated prostate cancer cell line migration and invasion. Nat Commun 11, 4498 (2020).

50. R. H. Straub, P. Harle, P. Sarzi-Puttini, and M. Cutolo. Tumor necrosis factor-neutralizing therapies improve altered hormone axes: an alternative mode of antiinflammatory action. Arthritis Rheum 54, 2039–2046 (2006).

51. D. W. Strand, D. N. Costa, F. Francis, W. A. Ricke, and C. G. Roehrborn. Targeting phenotypic heterogeneity in benign prostatic hyperplasia. Differentiation 96, 49–61 (2017).

52. T. Dang, and G. Y. Liou. Macrophage Cytokines Enhance Cell Proliferation of Normal Prostate Epithelial Cells through Activation of ERK and Akt. Sci Rep 8, 7718 (2018).

53. D. K. Smith, S. L. Hasanali, J. Wang, G. Kallifatidis, D. S. Morera, A. R. Jordan, M. K. Terris, Z. Klaassen, R. Bollag, V. B. Lokeshwar, and B. L. Lokeshwar. Promotion of epithelial hyperplasia by interleukin-8-CXCR axis in human prostate. Prostate 80, 938–949 (2020).

54. A. Bell-Cohn, D. J. Mazur, C. Hall, A. J. Schaeffer, and P. Thumbikat. Uropathogenic Escherichia coli-induced fibrosis, leading to lower urinary tract symptoms, is associated with type 2 cytokine signaling. Am J Physiol Renal Physiol 316, F682–F692 (2019).

55. J. B. Cheng, A. J. Sedgewick, A. I. Finnegan, P. Harirchian, J. Lee, S. Kwon, M. S. Fassett, J. Golovato, M. Gray, R. Ghadially, W. Liao, B. E. Perez White, T. M. Mauro, T. Mully, E. A. Kim, H. Sbitany, I. M. Neuhaus, R. C. Grekin, S. S. Yu, J. W. Gray, E. Purdom, R. Paus, C. J. Vaske, S. C. Benz, J. S. Song, and R. J. Cho. Transcriptional Programming of Normal and Inflamed Human Epidermis at Single-Cell Resolution. Cell Rep 25, 871–883 (2018).

56. N. MacRitchie, M. Frleta-Gilchrist, A. Sugiyama, T. Lawton, I. B. McInnes, and P. Maffia. Molecular imaging of inflammation - Current and emerging technologies for diagnosis and treatment. Pharmacol Ther 211, 107550 (2020).

57. M. Oleszowsky, W. Willinek, M. Marinova, and M. F. Seidel. Synovial inflammation analyzed by 3-T magnetic resonance imaging in etanercept-treated patients with rheumatoid arthritis indicates persistent disease activity despite of clinical remission: A pilot study. Mod Rheumatol 29, 441–446 (2019).

58. A. Dobin, C. A. Davis, F. Schlesinger, J. Drenkow, C. Zaleski, S. Jha, P. Batut, M. Chaisson, and T. R. Gingeras. STAR: ultrafast universal RNA-seq aligner. Bioinformatics 29, 15–21 (2013).

59. E. Z. Macosko, A. Basu, R. Satija, J. Nemesh, K. Shekhar, M. Goldman, I. Tirosh, A. R. Bialas, N. Kamitaki, E. M. Martersteck, J. J. Trombetta, D. A. Weitz, J. R. Sanes, A. K. Shalek, A. Regev, and S. A. McCarroll. Highly Parallel Genome-wide Expression Profiling of Individual Cells Using Nanoliter Droplets. Cell 161, 1202–1214 (2015).

60. R. Satija, J. A. Farrell, D. Gennert, A. F. Schier, and A. Regev. Spatial reconstruction of single-cell gene expression data. Nat Biotechnol 33, 495–502 (2015).

61. C. Hafemeister, and R. Satija. Normalization and variance stabilization of single-cell RNA-seq data using regularized negative binomial regression. Genome Biol 20, 296 (2019).

62. V. D. G. Blondel, Jean-Loup; Lambiotte, Renaud; Lefebvre, Etienne Fast unfolding of communities in large networks. Journal of Statistical Mechanics: Theory and Experiment 10, (2008).

63. L. Zappia, and A. Oshlack. Clustering trees: a visualization for evaluating clusterings at multiple resolutions. Gigascience 7, (2018).

64. Y. a. H. Benjamini, Y. Controlling the false discovery rate: a practical and powerful approach to multiple testing. Journal of the Royal Statistical Society Series B 57, 289– 300 (1995).

65. F. Wilcoxon. Individual Comparisons by Ranking Methods. Biometrics Bulletin 1, 80–83 (1945).

66. M. D. Robinson, D. J. McCarthy, and G. K. Smyth. edgeR: a Bioconductor package for differential expression analysis of digital gene expression data. Bioinformatics 26, 139–140 (2010).

67. D. J. McCarthy, Y. Chen, and G. K. Smyth. Differential expression analysis of multifactor RNA-Seq experiments with respect to biological variation. Nucleic Acids Res 40, 4288–4297 (2012).

68. M. Jiang, D. W. Strand, S. Fernandez, Y. He, Y. Yi, A. Birbach, Q. Qiu, J. Schmid, D. G. Tang, and S. W. Hayward. Functional remodeling of benign human prostatic tissues in vivo by spontaneously immortalized progenitor and intermediate cells. Stem Cells 28, 344–356 (2010).

69. O. E. Franco, M. Jiang, D. W. Strand, J. Peacock, S. Fernandez, R. S. Jackson, 2nd, M. P. Revelo, N. A. Bhowmick, and S. W. Hayward. Altered TGF-beta signaling in a subpopulation of human stromal cells promotes prostatic carcinogenesis. Cancer Res 71, 1272–1281 (2011).

70. R. E. Vickman, M. M. Broman, N. A. Lanman, O. E. Franco, P. A. G. Sudyanti, Y. Ni, Y. Ji, B. T. Helfand, J. Petkewicz, M. C. Paterakos, S. E. Crawford, T. L. Ratliff, and S. W. Hayward. Heterogeneity of human prostate carcinoma-associated fibroblasts implicates a role for subpopulations in myeloid cell recruitment. Prostate 80, 173–185 (2020).

71. A. F. Olumi, G. D. Grossfeld, S. W. Hayward, P. R. Carroll, T. D. Tlsty, and G. R. Cunha. Carcinoma-associated fibroblasts direct tumor progression of initiated human prostatic epithelium. Cancer Res 59, 5002–5011 (1999).

72. G. H. Henry, A. Malewska, D. B. Joseph, V. S. Malladi, J. Lee, J. Torrealba, R. J. Mauck, J. C. Gahan, G. V. Raj, C. G. Roehrborn, G. C. Hon, M. P. MacConmara, J. C. Reese, R. C. Hutchinson, C. M. Vezina, and D. W. Strand. A Cellular Anatomy of the Normal Adult Human Prostate and Prostatic Urethra. Cell Rep 25, 3530–3542 e3535 (2018).

