## Supplementary Materials for "TNF Blockade Reduces Prostatic Hyperplasia and Inflammation while Limiting BPH Diagnosis in Patients with Autoimmune Disease"

Figure S1

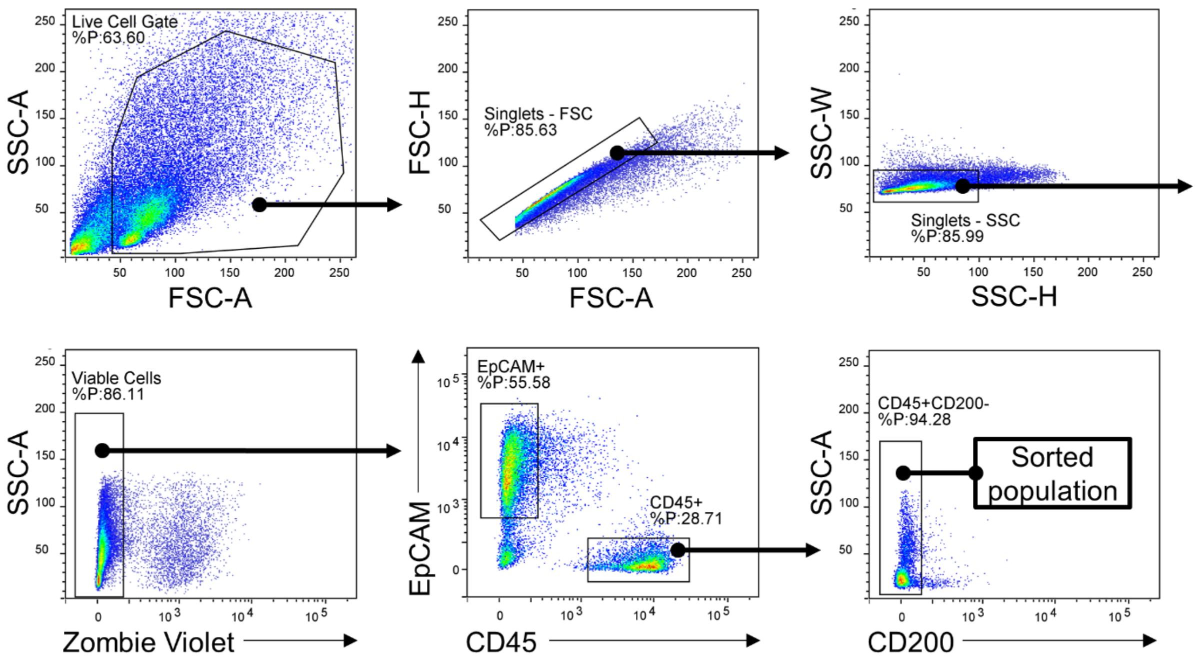


**Figure S1. Fluorescence activated cell sorting strategy for CD45+ scRNA-seq analysis.** Progressive gating strategy for isolation of CD45+ cells following enzymatic digestion of human prostate tissues, in order from the top left panel to the bottom right panel. Zombie violet was used for exclusion of dead cells and EpCAM and CD200 were used to exclude epithelial and endothelial cells (*72*), respectively.

Figure S2

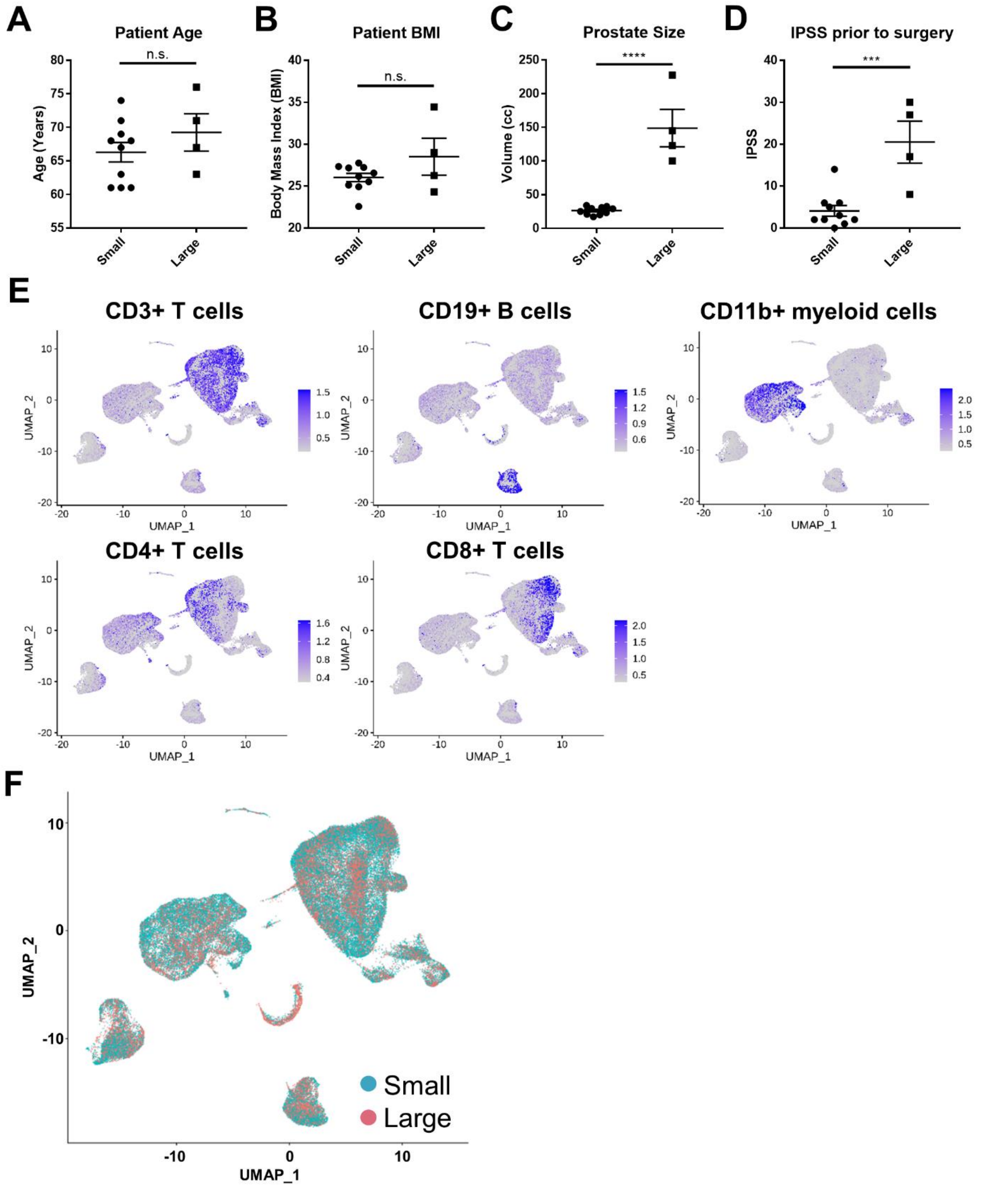


**Figure S2. Additional data relating to scRNA-seq data and patient characteristics. A)** Patient age, **B)** patient BMI, **C)** prostate size estimation by TRUS or CT scan, and **D)** IPSS prior to surgery is shown for ten patients with small prostates and four patients with large prostates. **E)** UMAP plot of 69,850 BPH associated cells, colored to highlight cells from small or large prostates, highlights the significant overlap between both populations. **F)** Feature plots highlighting protein expression of CD3 ($CD3^+$ T cells), CD4 ($CD4^+$ T cells), CD8 ($CD8^+$ T cells), CD19 ($CD19^+$ B cells), and $CD11b^+$ (myeloid cells). CITE-seq analysis was conducted on three small and three large samples.

Figure S3

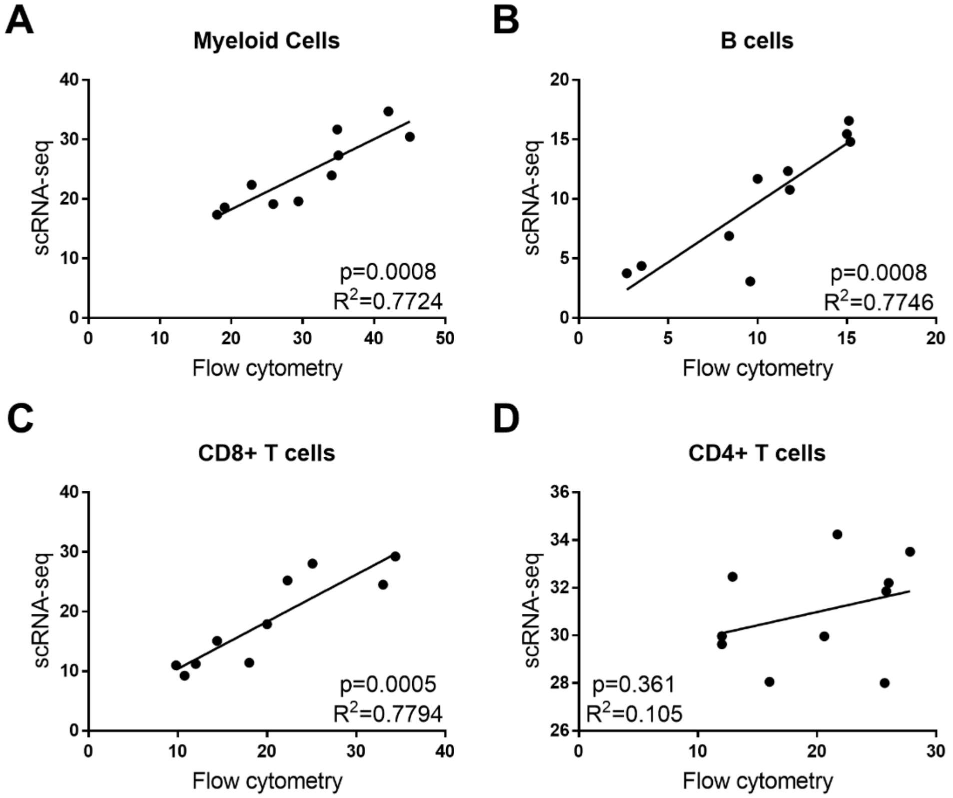


**Figure S3. Estimated proportions of immune cell types by scRNA-seq analysis represent immune cell proportions by flow cytometry.** Digested small (n=7) or large (n=3) prostate tissues were subjected to flow cytometry analysis after staining with antibodies against CD45, CD11b, CD19, CD4, and CD8. Sorted CD45+ cells were subjected to scRNA-seq analysis from the same digested prostate tissues. Linear regressions for the percentages of CD11b+ myeloid cells (A), CD19+ B cells (B), CD8+ T cells (C), and CD4+ T cells (D), are shown as the estimated percentages of total CD45+ cells via scRNA-seq *versus* flow cytometry. The respective p-values and $R^2$ values are indicated.

Figure S4

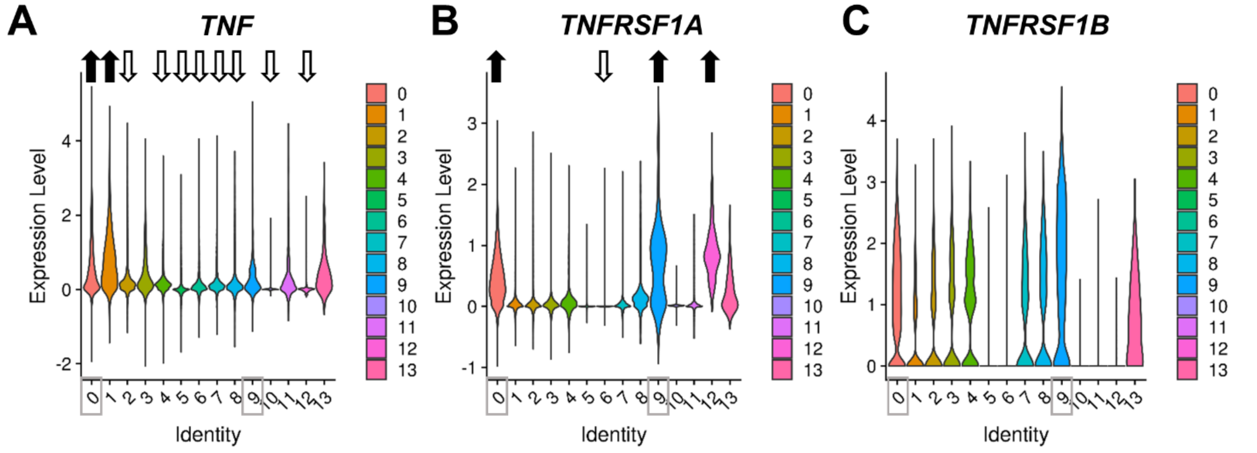


**Figure S4. Expression level of TNF, TNFRSF1A, and TNFRSF1B in BPH leukocytes.** Violin plots indicate the normalized gene expression level (log[CPM]) for **A)** *TNF*, **B)** *TNFRSF1A*, and **C)** *TNFRSF1B* in each of the clusters identified in the CD45+ scRNA-seq analysis. Arrows indicate clusters with significantly up- or down-regulated gene expression compared to all other clusters. Gray boxes highlight clusters 0 and 9 as the macrophage subpopulations.

Figure S5

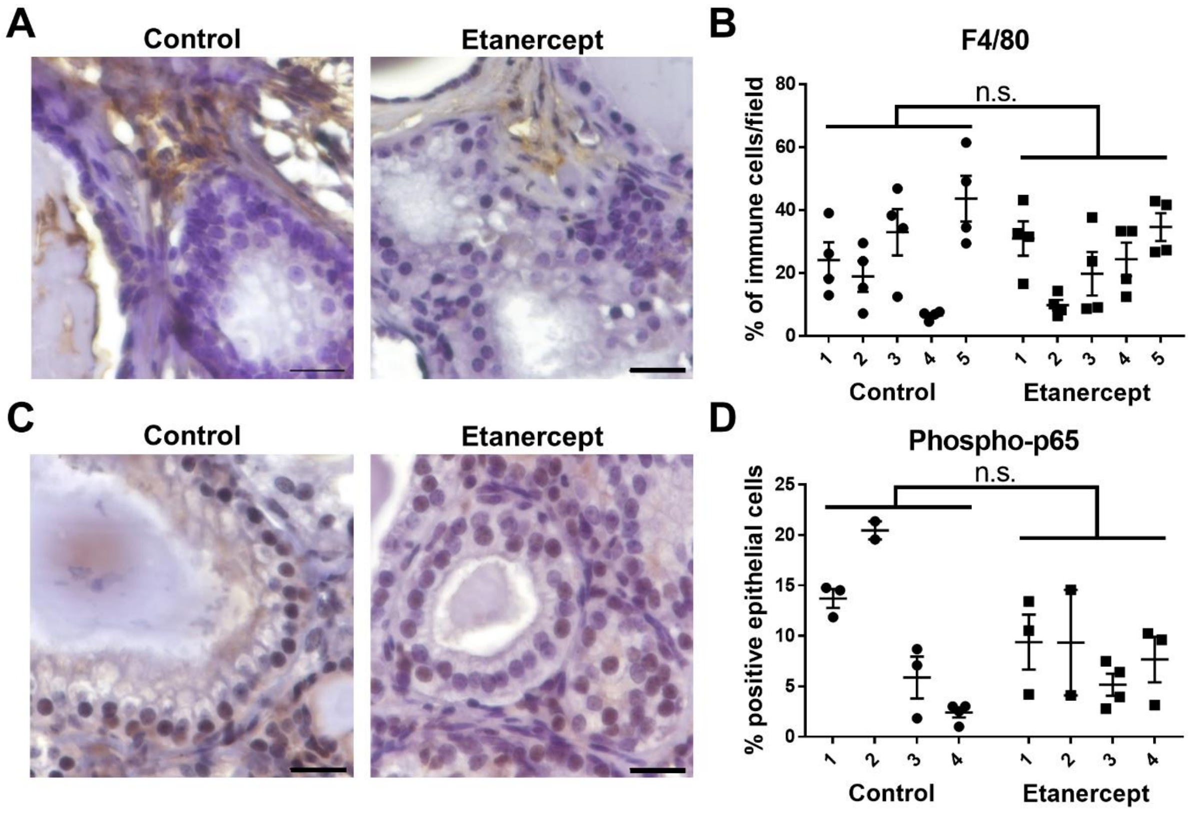


**Figure S5. TNFα-antagonist treatment in Pb-PRL mice does not significantly alter macrophage infiltration or epithelial NFκB activity. A)** Representative images of F4/80 staining in control or etanercept-treated mice. **B)** IHC staining counts for F4/80+ cells, represented as the portion of all immune cells in the field. **C)** Representative images of phospho-p65 staining in control or etanercept-treated mice. **D)** Data presented indicates the percentage of phospho-p65 positive epithelial cells counted per field of view after IHC staining. **B, D)** Data indicate the mean +/- SEM of the percent positive cells in at least three prostate tissue fields for each animal. Comparison of control and etanercept-treated groups for statistical purposes were conducted using a mixed effects model and the reported p-values indicate the significance of the treatment. Scale bars = 20µm. N.s.=not significant

Figure S6

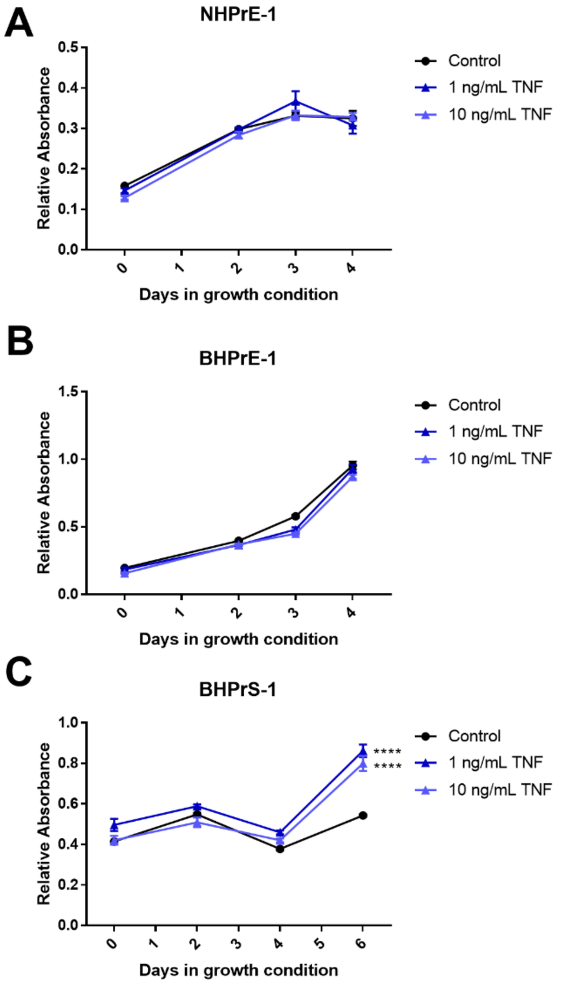


**Figure S6. TNF promotes stromal, but not epithelial, cell growth.** Crystal violet growth assays were performed in low serum conditions (0.5%) with indicated treatments. **A)** NHPrE-1, **B)** BHPrE-1, and **C)** BHPrS-1 were grown in the presence or absence of 1 or 10 ng/mL recombinant TNFα, followed by fixation for crystal violet growth assay at indicated times. Points indicate the mean +/- SEM of at least five technical replicates and graphs are representative of three independent experiments. Statistical analysis included a two-way ANOVA with Dunnett's multiple comparisons test, where **** represents $p<0.0001$ compared to control.

Figure S7

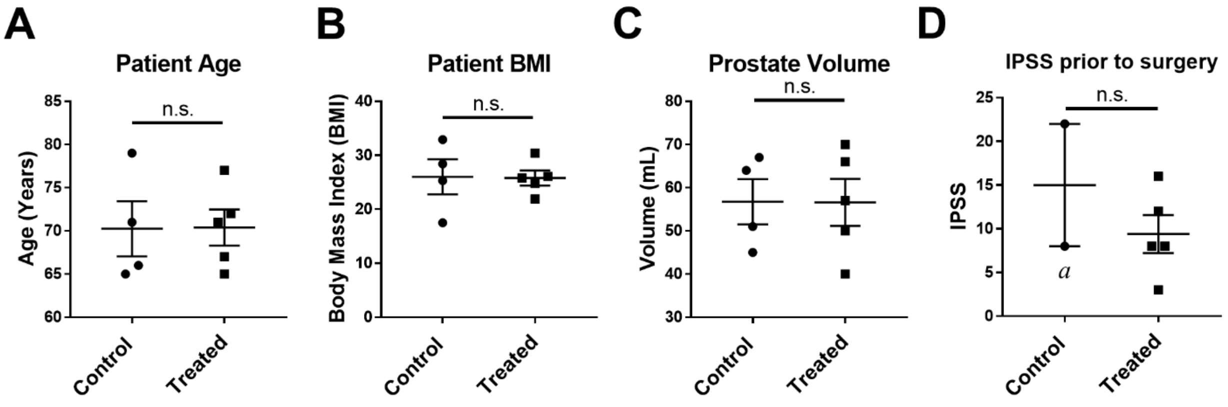


**Figure S7. Characteristics of TNFα-antagonist treated or control patients. A)** Patient age, **B)** patient BMI, **C)** prostate volume, and **D)** IPSS are shown for four control patients and five patients with large prostates, from which the relevant IHC data was generated. [a] two patients within the control group did not complete the IPSS questionnaire, thus have no values to report.

**Table S1. Proportion of men with BPH by autoimmune disease status.** Chi-square tests were utilized to compare the proportion of BPH diagnoses in men with an autoimmune (AI) condition *versus* men with no AI condition. One compares men with any diagnosis of an AI condition to men without, while the second comparison includes only men diagnosed with an AI condition prior to their clinical BPH diagnosis, as a subset of patients who may have been treated for their AI condition (9,274 AI patients were diagnosed with AI disease prior to a BPH diagnosis out of 10,769 total patients with AI disease). Subcategories of the AI disease population are provided. The bolded p-values indicate a significant difference in BPH prevalence compared to the "no autoimmune disease" group. [#] indicates a significantly higher BPH prevalence than the 20.3% reference, although the incidence rate in this subpopulation of RA patients is decreased from 38.0% prevalence in all RA patients.

| **Diagnosis of Men** | **Patients (%)** | **BPH Prevalence (N)** | **P-Value** |
|---|---|---|---|
| No Autoimmune Disease | 101,383 (90.4%) | 20.3% (20,586) | Reference |
| Autoimmune Disease (all) | 10,769 (9.6%) | 30.6% (3,294) | **<0.001** |
| • Psoriasis | 3,215 (2.9) | 24.9% | **<0.001** |
| • Rheumatoid Arthritis | 1,700 (1.5) | 38.0% | **<0.001** |
| • Ulcerative Colitis | 1,104 (1.0) | 30.2% | **<0.001** |
| • Type 1 Diabetes | 1,043 (0.9) | 32.0% | **<0.001** |
| • Crohn’s Disease | 971 (0.9) | 27.4% | **<0.001** |
| • Multiple Sclerosis | 375 (0.3) | 21.6% | 0.884 |
| • Celiac Disease | 373 (0.3) | 24.4% | 0.142 |
| • Hashimoto's Thyroiditis | 372 (0.3) | 22.9% | 0.463 |
| • Ankylosing Spondylitis | 287 (0.3) | 25.4% | 0.086 |
| • Lupus | 137 (0.1) | 30.7% | **0.007** |
| • Vasculitis | 47 | | |
| • Myasthenia Gravis | 32 | | |
| • Graves Disease | 3 | | |
| • Other | 2,613 (2.3) | | |
| Autoimmune Disease (prior to BPH diagnosis) | 9,274 of total 10,769 | 19.4% (1,799) | **0.037** |
| • Psoriasis | | 16.3% | **<0.001** |
| • Rheumatoid Arthritis | | 23.2%[#] | **0.007** |
| • Ulcerative Colitis | | 20.7% | 0.773 |
| • Type 1 Diabetes | | 19.4% | 0.521 |
| • Crohn’s Disease | | 19.4% | 0.521 |
| • Multiple Sclerosis | | 15.8% | **0.035** |
| • Celiac Disease | | 13.5% | **0.002** |
| • Hashimoto's Thyroiditis | | 11.4% | **<0.001** |
| • Ankylosing Spondylitis | | 18.0% | 0.357 |
| • Lupus | | 16.7% | 0.334 |

**Table S2. Median Days for Progression to Surgery for BPH Patients.** For n=1,685 patients, data indicate the median number of days between BPH diagnosis and surgery. The median days for progression to surgery is shown between men taking methotrexate *versus* men not taking methotrexate, as well as for men taking TNFα-antagonists *versus* men not taking TNFα-antagonists. Analysis was completed using the Mann-Whitney U test and p-values are shown. Two patients that progressed to surgery were taking both methotrexate and TNFα-antagonists. [a]Patients taking methotrexate compared to those not taking methotrexate (n=1670); [b]Patients taking TNFα-antagonists compared to those not taking TNFα-antagonists (n=1678)

| **BPH Patient Group** | **Median Days** | **P-Value** |
|---|---|---|
| All BPH patients who progressed to surgery (n=1,685) | 616 | |
| • Not taking methotrexate or TNFα-antagonists (n=1,665) | 613 | |
| • Taking methotrexate (n=15) | 728 | 0.75[a] |
| • Taking TNFα-antagonists (n=7) | 1,115 | 0.15[b] |

**Table S3. Most significantly altered pathways in CD45+ cells from large versus small human BPH tissues.** Ingenuity Pathway Analysis (Qiagen) was conducted using the differentially expressed genes between large *versus* small CD45+ scRNA-seq samples. Gray highlighted rows indicate pathways related to AI diseases.

| **Ingenuity Canonical Pathways** | **-Log(p-value)** | **Ratio** |
|---|---|---|
| Communication between Innate and Adaptive Immune Cells | 11.5 | 0.177 |
| Hematopoiesis from Pluripotent Stem Cells | 10 | 0.245 |
| EIF2 Signaling | 7.94 | 0.0893 |
| NRF2-mediated Oxidative Stress Response | 7.64 | 0.0952 |
| Systemic Lupus Erythematosus In B Cell Signaling Pathway | 7.13 | 0.0764 |
| Atherosclerosis Signaling | 6.91 | 0.111 |
| B Cell Receptor Signaling | 6.29 | 0.0865 |
| Role of Macrophages, Fibroblasts and Endothelial Cells in Rheumatoid Arthritis | 6.21 | 0.0673 |
| IL-10 Signaling | 6.17 | 0.145 |
| IL-6 Signaling | 6.12 | 0.104 |
| Dendritic Cell Maturation | 5.64 | 0.082 |
| Primary Immunodeficiency Signaling | 5.38 | 0.16 |
| Systemic Lupus Erythematosus Signaling | 5.08 | 0.0699 |
| Granulocyte Adhesion and Diapedesis | 5.04 | 0.0778 |
| Cholecystokinin/Gastrin-mediated Signaling | 4.78 | 0.0924 |
| Graft-versus-Host Disease Signaling | 4.5 | 0.146 |
| Autoimmune Thyroid Disease Signaling | 4.44 | 0.143 |
| Allograft Rejection Signaling | 4.44 | 0.105 |

**Table S4. Run metrics for scRNA-seq of BPH associated CD45+ leukocytes.** Table includes the run metrics from each scRNA-seq sample, including 10 small and 4 large samples. The metrics indicate high quality data and that the average number of cells captured approached the intended number of 5000 cells. The final row indicates the average indicated metric for all samples.

| **Sample** | **Number of Reads** | **% Q30 Bases in Read** | **% Reads Mapped** | **Estimated Number of Cells** | **Mean Reads per cell** | **Median Genes per cell** |
|---|---|---|---|---|---|---|
| **003_small** | 437,303,375 | 91.3 | 96.2 | 6,388 | 68,457 | 1692 |
| **004_small** | 266,767,655 | 91.7 | 96.7 | 4,163 | 64,080 | 1599 |
| **006_small** | 258,138,447 | 91.5 | 96.9 | 6,427 | 40,164 | 1628 |
| **007_small** | 765,500,750 | 90.8 | 96.0 | 4,427 | 172,916 | 1958 |
| **008_small** | 552,744,091 | 93.0 | 98.1 | 5,053 | 109,389 | 1652 |
| **009_small** | 361,879,675 | 93.3 | 96.7 | 3,659 | 98,901 | 1740 |
| **010_small** | 245,628,503 | 91.6 | 96.3 | 3,750 | 65,500 | 1576 |
| **012_large** | 297,618,482 | 90.2 | 96.7 | 5,826 | 51,084 | 1526 |
| **013_small** | 393,520,430 | 92.6 | 96.7 | 5,230 | 75,242 | 1476 |
| **1144_small** | 394,479,925 | 90.0 | 97.0 | 6,010 | 65,637 | 1662 |
| **1157_large** | 303,893,852 | 90.4 | 96.8 | 6,816 | 44,585 | 1525 |
| **1195_large** | 455,027,712 | 92.4 | 96.6 | 5,377 | 84,624 | 1591 |
| **1196_small** | 224,350,221 | 92.7 | 96.8 | 4,554 | 49,264 | 1489 |
| **766_large** | 368,324,247 | 90.8 | 95.6 | 4,076 | 90,364 | 1335 |
| **AVERAGE:** | **380,369,812** | **91.6** | **96.7** | **5,125** | **77,158** | **1,604** |

**Table S5. Antibodies used for IHC in human and mouse tissues.** Table indicates the antibody target, clone (if monoclonal), manufacturer product number, and dilution used for all antibodies used for IHC studies.

| Tissue Sample | Antibody Target | Clone | Species | Isotype | Company | Ref. # | IHC Dilution |
|---|---|---|---|---|---|---|---|
| Human prostate | phospho-NFkB p65 (S276) | polyclonal | Rabbit | IgG | Abcam | ab194726 | 1:100 |
| Human prostate | Ki67 | polyclonal | Rabbit | IgG | Abcam | ab15580 | 1:50 |
| Human prostate | CD68 | EPR20545 | Rabbit | IgG | Abcam | ab213363 | 1:100 |
| NOD prostate | F4/80 | polyclonal | Rabbit | IgG | Abcam | ab100790 | 1:100 |
| NOD prostate | Ki67 | polyclonal | Rabbit | IgG | Abcam | ab15580 | 1:75 |
| NOD prostate | phospho-NFkB p65 (S276) | polyclonal | Rabbit | IgG | Abcam | ab194726 | 1:100 |
| PB-PRL prostate | F4/80 | polyclonal | Rabbit | IgG | Abcam | ab100790 | 1:100 |
| PB-PRL prostate | Ki67 | polyclonal | Rabbit | IgG | Abcam | ab15580 | 1:200 |
| PB-PRL prostate | phospho-NFkB p65 (S276) | polyclonal | Rabbit | IgG | Abcam | ab194726 | 1:100 |
